# Targeting Interferon-Driven Inflammatory Memory Prevents Epigenetic Evolution of Cancer Immunotherapy Resistance

**DOI:** 10.1101/2024.08.13.607862

**Authors:** Jingya Qiu, Darwin Ye, Xinyi E. Chen, Nathan Dangle, Benjamin Yoshor, Thomas Zhang, Yue Shao, Vamshidhar C. Nallamala, Shangshang Wang, Diqiu Ren, Yuanming Xu, Jie Chen, Nancy R. Zhang, Junwei Shi, Roger A. Greenberg, Andy J. Minn

**Author notes:** Corresponding author and lead contact: Andy J. Minn, MD, PhD, 421 Curie Blvd BRB II/III, Room 510, Philadelphia, PA 19104. Clinical Pharmacology & Translational Sciences, Pfizer Oncology Unit, Pfizer-La Jolla, San Diego, CA 92121. These authors contributed equally (Jingya Qiu, Darwin Ye).

## Abstract

Acquired resistance is a growing obstacle to durable responses after cancer immune checkpoint blockade (ICB). The mechanisms by which heterogeneous tumors evolve under immunotherapy pressure and strategies targeting key populations to prevent relapse are poorly understood. We show that chronic interferon (IFN) enables a subpopulation of cancer cells to acquire inflammatory memory and express memory ISGs, a subset of IFN-stimulated genes enriched for immune evasion properties, leading to subclonal epigenetic evolution of ICB-resistant states. Inflammatory memory is epigenetically encoded through chronic virus mimicry – feedforward MDA5 signaling likely activated by endogenous retroelements. While JAK inhibitors can improve ICB response, combining them with TBK1 inhibitors collapses this feedforward mechanism, erasing inflammatory memory and preventing differentiation into resistance states. Across human cancers, small subpopulations of memory ISG-expressing cells are prevalent and coupled to T cell exhaustion, suggesting inflammatory memory may be a common mechanism of acquired resistance targetable by JAK plus TBK1 inhibition.

## INTRODUCTION

Cancer immunotherapy seeks to coordinate an immune response against tumors by activating innate and/or adaptive immunity^1^. Immune checkpoint blockade (ICB) agents that include anti-PD1 and anti-PDL1 accomplishes this by preventing the engagement of inhibitory receptor signaling on adaptive immune cells such as CD8 T cells, or on innate immune cells such as myeloid and dendritic cells^2–4^. Productive cellular interactions involving these immune cell types together with cancer cells can be critical for priming an effective anti-tumor immune response and ICB efficacy^5^. Consistent with this, features that indicate the presence of T cells, their recruitment and potential interaction with myeloid and dendritic cells, and their likelihood of recognizing tumor antigens on cancer cells can all be associated with patients who respond to ICB^6,7^. Many of these features are positively influenced by interferon (IFN), including type one IFN (IFN-I) or type two IFN (IFNG) signaling, explaining why expression of IFN-stimulated genes (ISGs) associates with immunotherapy efficacy.

Despite the increasing success of immunotherapies like ICB across many different cancer types, relapse after initial response, also referred to as acquired resistance, has become an increasing barrier to durable response and long-term survival^8^. Unexpectedly, IFN signaling and ISGs can also be associated with immunotherapy failure and acquired resistance. For example, tumors from patients with non-small cell lung cancer (NSCLC) that relapse after anti-PD1 often exhibit increased levels of a subset of ISGs and gene signatures for immune dysfunction^9^. This subset of ISGs and features of immune dysfunction have also been observed in mouse tumors that relapsed after immunotherapy^10,11^, and high expression of similar subsets of ISGs can predict poor ICB response in melanoma^12^ and non-durable response to CD19 CAR-T cells^13^. Accordingly, preventing IFN signaling can sensitize mouse tumors to ICB, and genes involved in the IFN pathway are a dominant category for immune evasion genes in genome-wide *in vivo* CRISPR screens^14^. Furthermore, we recently showed in a clinical trial for metastatic NSCLC that addition of a JAK inhibitor (JAKi) to interfere with IFN and inflammatory cytokine signaling holds promise as a therapeutic approach to improve anti-PD1^15^, results independently seen in another clinical trial for refractory Hodgkin lymphoma^16^. Here, the combination of JAKi with anti-PD1 demonstrated high clinical response rates and improved CD8 T cell and myeloid cell function in NSCLC and lymphoma patients, respectively. However, we also observed a subset of NSCLC patients who appeared refractory to JAK inhibition and failed to respond, highlighting an opportunity to understand the mechanism for this effect and to improve upon JAKi. Together, these findings suggest that besides initial coordinating events between immune cells and cancer cells important for immune priming, IFN signaling and a specific subset of ISGs function in a different biological and clinical context to promote immune suppression, possibly contributing to mechanisms of acquired immunotherapy resistance. Building on the use of JAKi to target ICB resistance related to IFN and inflammatory signaling holds promise as a therapeutic strategy.

One context that captures the opposing roles of IFN signaling in immune regulation is acute versus chronic viral infections^17^. Here, chronic LCMV infection in mice^18,19^, SIV/HIV infection in primates and humans^20,21^, and chronic viral hepatitis^22^ are all associated with persistent IFN-I signaling that promotes immune suppression. This IFN-driven immunomodulation is thought to be an adaptive response to decrease the risk of immune-mediated pathology^17^. Part of mitigating risk of inflammation-induced pathology is the ability not just to constrain inflammation but also to promote tissue regeneration and restore homeostasis. Such an adaptive response to inflammation has been described in epithelial and other normal tissue types as inflammatory memory and may be particularly relevant for stem and progenitor cells from these tissues^23,24^. For cancer, we have recently described how ICB resistant tumors acquire IFN-driven epigenetic features resembling inflammatory memory^25^. Here, chronic IFNG stimulation of cancer cells leads to ICB resistance and the development of poised chromatin to enhance tonic expression of ISGs that augment IFN-I production. This IFN-I secretion by cancer cells may be a source of chronic IFN-I that sustains not only ICB resistance of cancer cells but also directly contributes to T cell exhaustion in the tumor microenvironment^25,26^. How IFN-related inflammatory memory in cancer cells contributes to immune dysfunction and whether this process relates to pathways activated during chronic virus infections are poorly understood.

Within the tumor microenvironment, clonal variation found in cancer cells is thought to be an important factor in cancer therapy resistance^27^. For tyrosine kinase inhibitors (TKIs) that target driver oncogenes, this heterogeneity can result in subclones with somatic mutations that lead to bypass signaling^28^. For ICB resistance, somatic mutations in genes like B2M critical for MHC-I expression, JAK1/2 important in IFNG signaling, or STK11/LBK1 have been described^8,29^. However, such genetic mechanisms seem to account for only a small fraction of ICB resistance^8^. In the case of TKI resistance, non-genetic mechanisms appear involved in how multiple clonal fates can emerge from seemingly homogenous cancer cells^30^. These findings point toward the concept that epigenetic features might vary and evolve among cancer cell subclones and be an important source of therapy resistance. The extent to which epigenetically distinct subclones contribute to ICB resistance, the mechanisms involved, and how these non-genetic resistance pathways can be targeted are outstanding questions.

In this study, we investigate how inflammatory memory from chronic IFNG signaling leads to the development of an ICB-resistant epigenetic state that originates from a subpopulation of cancer cells found across human cancers. This resistant epigenetic state is sustained by a feed-forward chronic antiviral signaling mechanism. This mechanism can be collapsed by the combination of JAK and TBK1 inhibitors, highlighting how inflammatory memory responsible for an epigenetic mechanism of acquired immunotherapy resistance can potentially be targeted.

## RESULTS

### Chronic IFN signaling and immunotherapy resistance are linked to memory ISGs with epigenetic features of inflammatory memory

We previously demonstrated that cancer cells that relapsed after ICB can possess epigenetic features of inflammatory memory^25^. Persistent IFN signaling, particularly through IFNG, is an important mediator of this effect as chronic IFNG stimulation *in vitro* is sufficient to render cancer cells resistant to ICB and initiate epigenetic reprogramming^11^. To characterize how the acquisition of IFNG-driven resistance and epigenetic features of inflammatory memory alter the transcriptional response to IFNG, parental B16-F10 (B16) melanoma cells, B16 cells rendered ICB resistant by chronic IFNG treatment *in vitro* (B16*y*), and B16 cells from tumors that relapsed after ICB (Res 499)^10^ were stimulated with IFNG for 6 hours *in vitro* **(Figure 1A)**. After washout, cells were collected for transcriptome and chromatin accessibility profiling over a 144-hour period **(Figure 1B)**. We empirically identified ISGs by hierarchically clustering B16 and Res 499 gene expression across time points, revealing three classes of ISGs with disparate temporal patterns **(Figure S1A-C)**. “Resolved ISGs” (ISG-resolved) were induced by IFNG after 6 hours and promptly returned to unstimulated levels after washout (**Figure 1C**). “Wave 2” ISGs (ISG-wave2) had similar induction kinetics, resolved at 12 hours post-washout, and displayed a second wave of transcription that peaked at 72 hours (**Figure S1C**). Neither group of ISGs differed between ICB-resistant Res 499 and parental B16 cells. In contrast, a third group of ISGs called “Memory ISGs” (ISG-mem) exhibited different kinetics in Res 499, and to a lesser extent in B16*y*, compared to parental B16 (**Figure 1C-D**). Specifically, this subset of 45 ISGs showed greater induction at 6 hours and persistent expression even at 72-144 hours post-washout. Thus, a select class of ISGs in ICB-resistant cancer cells respond to IFNG with enhanced induction and persistence.

**Figure 1.**
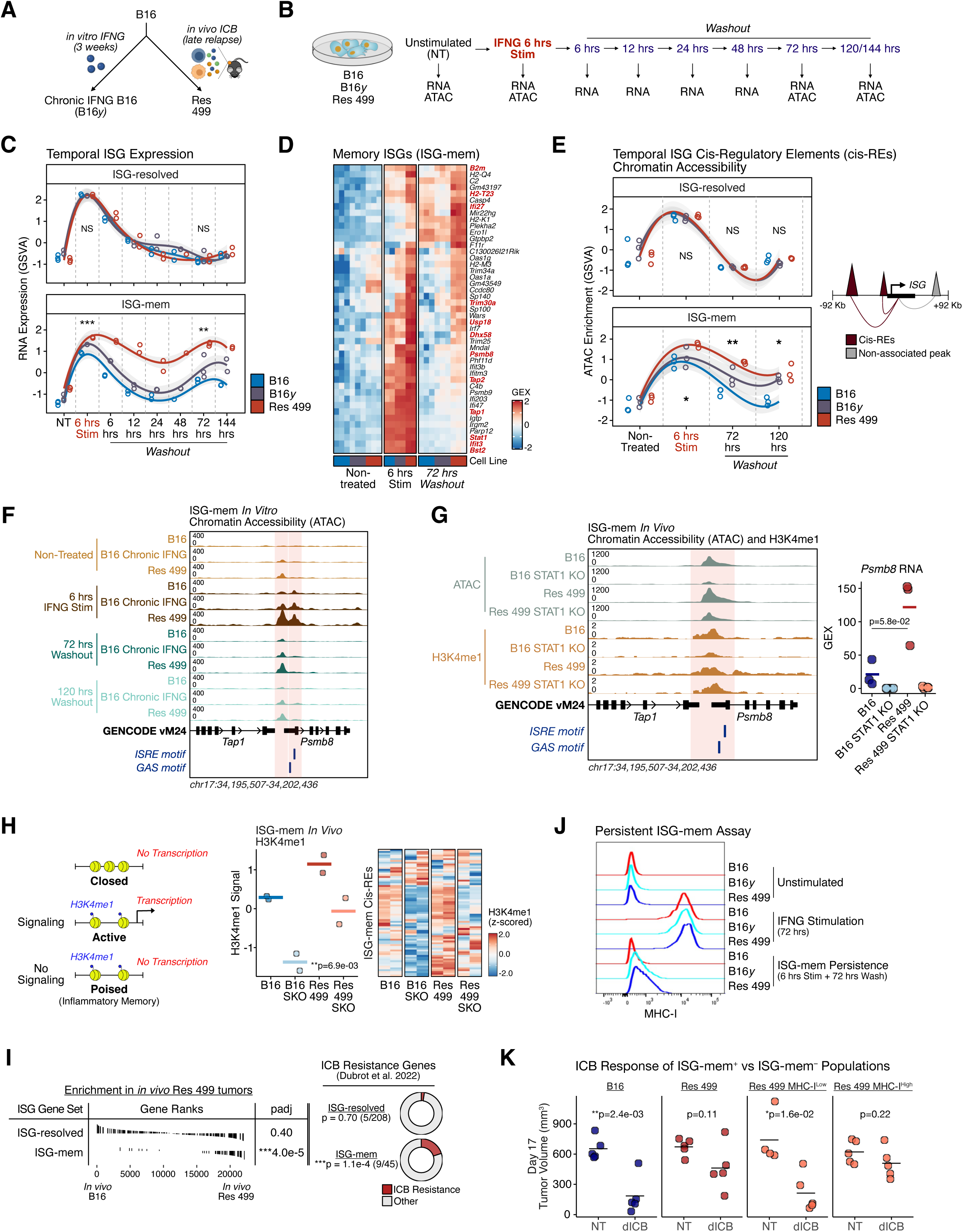
Acquired immunotherapy resistance and chronic IFNG signaling results in inflammatory memory and persistence of memory ISGs enriched in immune evasion genes. **A)** B16 melanoma cells rendered resistant from *in vitro* IFNG stimulation for 3 weeks (B16*y*) or isolated from an *in vivo* tumor after ICB relapse (Res 499). **B)** Schema of IFN stimulation and washout experiment to examine ISG chromatin and transcriptional dynamics. **C)** Summary gene set enrichment score (GSVA) of ISG-resolved and ISG-mem in the indicated cancer cell line across sampled time points (n=2-3 mice per condition and time point). A smoothed spline regression line with 95% confidence intervals is fit for each cancer cell line. Asterisks denote whether gene expression significantly differs between B16 and Res 499 at the indicated time point. **D)** Scaled gene expression in the indicated cancer cell line for all ISG-mem genes summarized in Figure 1C (bottom). Resistance genes are highlighted in red. **E)** Chromatin accessibility score at putative cis-regulatory elements (cis-REs) linked to ISG-resolved or ISG-mem signatures (right schema) assessed by ATAC GSVA enrichment score in the indicated cancer cell line and time point (left; n=3 mice per condition and time point). A smoothed spline regression line with 95% confidence intervals is fit for each cancer cell line. Asterisks denote whether chromatin accessibility significantly differs between B16 and Res 499 at the indicated time point. **F)** Representative ATAC-seq tracks for the indicated *in vitro* condition at an ISG-mem loci. GENCODE vM24 gene annotations and instances of indicated TF archetype motifs (AC0672, AC0668) are shown. **G)** Representative ATAC-seq or H3K4me1 CUT&RUN tracks at an ISG-mem loci (left) along with gene expression values (right) for the indicated B16 or Res 499 cells with or without STAT1 KO sorted from *in vivo* tumors. **H)** Schema of chromatin features of inflammatory memory (left), and summary and heatmap of H3K4me1 GSVA enrichment scores for ISG-mem cis-REs from B16 or Res 499 cells with or without STAT1 KO (SKO) sorted from *in vivo* tumors (middle and right). P-value was determined by a one-way ANOVA test. **I)** Gene set enrichment analysis (GSEA) results for ISG-resolved or ISG-mem on genes ranked by differential expression between Res 499 and B16 *in* vivo tumors (left). Over-representation analysis results for enrichment of Dubrot et al. ICB-resistance genes in each ISG subset (right). P-values determined by Fisher’s exact test. **J)** Persistent ISG-mem assay measured by MHC-I, a memory ISG. Shown is a representative flow cytometry histogram of MHC-I in B16, B16*y*, or Res 499 cells after the indicated *in vitro* treatment. **K)** Day 17 tumor volumes for B16, Res 499, or Res 499 sorted for low or high levels of MHC-I after *in vitro* IFNG stimulation plus washout. Mice were either non-treated or treated with anti-CTLA4 +/- anti-PDL1 (ICB). Unless otherwise indicated, P-values for two sample comparisons were determined by a two-sided t-test. *p < 0.05; **p<0.01; ***p<0.001.

We next assessed if the altered transcriptional dynamics of ISG-mem are associated with distinct epigenetic traits. We used a previously described regression model to identify peak-to-gene links, or putative cis-regulatory elements (cis-REs), within a 184 Kb cis-regulatory window for each ISG^25,31^. Consistent with the transcriptional dynamics, chromatin accessibility at putative cis-REs of ISG-mem remained elevated for up to 120 hours after *in vitro* IFNG stimulation in B16*y* and Res 499 cells, but not in B16 cells **(Figure 1E-F)**. Examination of cancer cells sorted from *in vivo* tumors confirmed that ISG-mem cis-REs, but not ISG-resolved cis-REs, also exhibited increased chromatin accessibility in Res 499 tumors **(Figure 1G, green tracks; Figure S1D-E)**. This increased chromatin accessibility of ISG-mem *in vivo* was also accompanied by increased H3K4me1, a histone modification associated with inflammatory memory that marks “poised” chromatin, or chromatin that remains open even in the absence of inflammatory signaling and gene transcription^23^ **(Figure 1H)**. To confirm that ISG-mem cis-REs represent poised chromatin, we knocked out STAT1 in Res 499 and B16 tumors to prevent IFN signaling and gene transcription. This revealed that cis-REs for many ISG-mem genes, like *Tap1* and *Psmb8*, indeed maintained enhanced chromatin accessibility and H3K4me1 levels *in vivo* even after STAT1 deletion in Res 499 compared to B16 tumors **(Figure 1G-H; Figure S1D)**, while no discernible effects were seen with cis-REs for ISG-resolved genes **(Figure S1E)**. Together, these data suggest that chronic IFNG stimulation and ICB-resistance of cancer cells lead to the acquisition of ISG-mem with greater induction and persistent expression days after termination of IFNG stimulation. These enhanced transcriptional dynamics are accompanied by acquisition of poised chromatin characteristic of inflammatory memory.

### Memory ISGs are enriched in regulators of immune evasion, antiviral signaling, and inflammation

The acquisition of persistent ISG-mem during the development of resistance after chronic IFNG stimulation or ICB relapse suggests that this subset of ISGs might contribute to immune evasion. Indeed, a large fraction of immune evasion genes identified by genome-wide *in vivo* CRISPR screening of multiple mouse tumor models treated with immunotherapy^14^ are ISGs in B16 and Res 499 cells **(Figure S1F)**, and 20% (9 out of 45) of ISG-mem genes are immune evasion genes **(Figure 1I, right donut plots; Figure S1G)**. In contrast, only 2% of ISG-resolved and 3% of ISG-wave2 overlap with this genome-wide collection of immune evasion genes. ISG-mem that overlap with immune evasion genes **(Figure 1D, red labels)** includes *H2-T23*, which is a non-classical MHC-I ligand for the NKG2A inhibitory receptor expressed on T cells and NK cells^14,32^, and *Usp18*, which is a ubiquitin-specific protease that modulates ISG expression and inhibits immunogenic cell death of cancer cells^33^. Other ISG-mem genes are involved in RNA sensing and antiviral signaling (e.g., *Dhx58* and *Trim25*)^34,35^, the epigenetic or post-translational control of pathological inflammation (e.g., *Sp140* and *Irgm2*)^36,37^, or are transcription factors important for IFN and antiviral gene expression (e.g., *Stat1* and *Irf7*)^35^. Importantly, ISG-mem, but not ISG-resolved or ISG-wave2, were significantly upregulated *in vivo* from ICB-resistant Res 499 compared to parental B16 tumors **(Figure 1I, left GSEA plot; Figure S1G)**. These findings suggest that ISG-mem enriches for at least three main functional categories: immune evasion, antiviral signaling, and modulation of inflammatory responses. Thus, the epigenetic features of inflammatory memory acquired by cancer cells may serve to control nucleic acid sensing, inflammatory sequelae, and ICB resistance.

To confirm that the acquisition of ISG-mem is coupled to ICB resistance, we identified cells that have acquired ISG-mem by examining cells with surface expression of MHC-I, which is an ISG-mem gene **(Figure 1D)**, that persists after IFNG stimulation. Accordingly, B16 cells increased MHC-I after 6 hours of *in vitro* IFNG stimulation before promptly returning to baseline **(Figure 1J, Figure S1H)**. In contrast, Res 499 cells and B16*y* cells rendered resistant by chronic IFNG had persistently elevated MHC-I for 72-120 hours in a subpopulation of cells after IFNG stimulation, matching ISG-mem transcriptional dynamics. To establish that Res 499 cells that acquired persistent MHC-I were indeed ICB-resistant, we isolated the Res 499 populations with and without persistent MHC-I after IFNG stimulation by cell sorting **(Figure S1I)**. Implantation of MHC-I^HIGH^ or MHC-I^LOW^ cells into mice followed by treatment with anti-PDL1 plus anti-CTLA4 revealed that MHC-I^HIGH^ tumors were more ICB resistant than MHC-I^LOW^ tumors **(Figure 1K)**. Thus, ISG-mem are enriched in immune evasion genes, and cancer cells capable of acquiring ISG-mem expression may contribute to ICB resistance.

### Chronic antiviral signaling involving MDA5 and endogenous retroelements maintain inflammatory memory and resistance

Because transcription and chromatin accessibility of ISG-mem are maintained long after the termination of IFNG stimulation, we surmised that resistant cancer cells may have acquired additional pathways to activate IFN signaling in a cell autonomous manner, reminiscent of a feedforward circuit. As noted, ISG-mem were also enriched in antiviral signaling genes, particularly those related to RIG-I-like receptors (RLRs) such as MDA5 and RIG-I that sense cytoplasmic RNAs^38^. These RNA ligands are thought to include endogenous retroelements (EREs) like endogenous retroviruses (ERVs), long interspersed nuclear elements (LINEs), and short interspersed nuclear elements (SINEs). Thus, we quantified EREs from tumors derived from B16, B16 cells stimulated *in vitro* for either 6 hours (acute) or 3.5 weeks (chronic) prior to implantation (B16*y*), and Res 499. This analysis revealed a small set of EREs, predominantly ERVs, exhibited elevated expression accompanied by increased chromatin accessibility and H3K4me3-marked transcriptional start sites, specifically in cancer cells from B16*y* and ICB-relapsed Res 499 tumors **(Figure 2A-B)**. Consistent with these findings, immunostaining for cytoplasmic double-stranded RNA (dsRNA) revealed significantly higher *ex vivo* dsRNA levels in cancer cells derived from Res 499 or B16*y* tumors compared to B16 tumors **(Figure 2C)**. The coupling of ERE and dsRNA levels to the acquisition of ISG-mem was also observed in KP (Kras^G12D^; Trp53^flox/flox^) mouse lung cancer cells either rendered resistant after chronic IFNG stimulation or derived from ICB-relapsed tumors **(Figure 2D-F; Figure S2A-G)**. Together, these data suggest that in multiple mouse cancer models of ICB resistance, acquisition of transcriptional and epigenetic features of ISG-mem is accompanied by de-repression of EREs.

**Figure 2.**
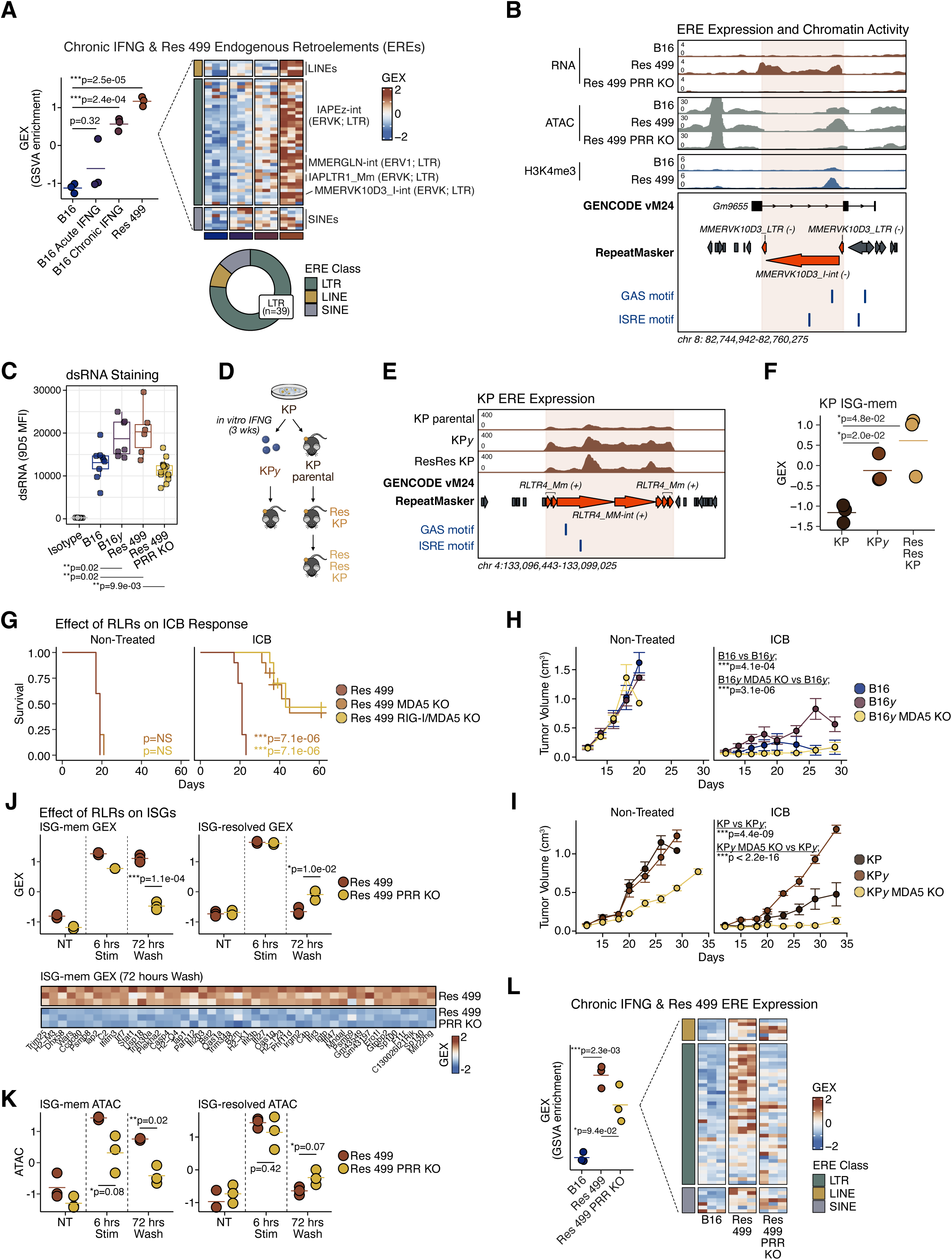
Acquired resistance and inflammatory memory are associated with chronic virus mimicry through endogenous retroelements and MDA5-dependency. **A)** Summary GSVA enrichment scores and heatmap for expression of endogenous retroelements (EREs) significantly increased in both B16*y* and Res 499 cells sorted from *in vivo* tumors (n=3 mice per condition). The ERE class composition is indicated in a donut plot (bottom). Heatmap shows individual biological replicates in the same order as the summary plot. **B)** Representative RNA-seq, ATAC-seq, and H3K4me3 CUT&RUN tracks at an ERE loci identified in (A) for B16 or Res 499 cancer cells with or without MDA5/RIG-I knockout (PRR KO) sorted from *in vivo* tumors. GENCODE vM24 and RepeatMasker annotations and instances of indicated TF archetype motifs (AC0668, AC0672) are shown. **C)** Mean fluorescence intensity (MFI) of dsRNA levels using the 9D5 antibody in the indicated tumors (n=6-9 mice per condition). **D)** Schema depicting generation of ICB-resistant KP lung cancer cells by *in vitro* chronic IFNG stimulation for 3 weeks (KP*y*) or after serial ICB relapse of *in vivo* KP tumors (Res KP, ResRes KP). **E)** Representative RNA-seq tracks at an ERE loci de-repressed after ICB relapse or chronic IFNG stimulation for the indicated KP cells sorted from *in vivo* tumors. **F)** Summary GSVA enrichment score for ISG-mem expression in the indicated KP cells sorted from *in vivo* tumors (n=3 per condition). **G)** Survival of mice with Res 499 parental, MDA5 knockout (KO), or MDA5/RIG-I double KO tumors either non-treated (n=5 per condition) or treated with anti-PDL1 + anti-CTLA4 (ICB) (n=10 per condition). **H)** Tumor growth of mice with B16, B16*y*, or B16*y* MDA5 knockout (KO) tumors either non-treated (n=5 per condition) or treated with anti-CTLA4 + anti-PDL1 (ICB) (n=10 per condition). **I)** Tumor growth of mice with KP, KP*y*, or KP*y* MDA5 KO tumors either non-treated (n=5 per condition) or treated with anti-CTLA4 + anti-PD1/anti-PDL1 + CD40 agonist (ICB) (n=10 per condition). **J)** Summary GSVA enrichment score for expression of memory and resolved ISGs in Res 499 cells with and without MDA5/RIG-I double KO (PRR KO) after 6 hours IFNG stimulation and at 72 hours after washout (schema shown in Fig. 1B). Expression of each ISG-mem at the 72 hour washout time point for individual biological replicates is visualized in a heatmap (bottom). **K)** Summary ATAC GSVA enrichment score for chromatin accessibility of memory and resolved ISG cis-REs. **L)** Expression of EREs from (A) in cancer cells sorted from the indicated *in vivo* tumor are shown in the heatmap as individual replicates, along with a plot of summary GSVA enrichment scores (left). P-values for two sample comparisons were determined by a two-sided t-test. P-values for tumor growth were determined by a mixed-effect regression model, and for survival by a log-rank test. Error bars represent standard errors.

Elevated EREs and dsRNA suggest that ICB-resistant cancer cells undergo chronic antiviral signaling reminiscent of a persistent viral infection and may rely on “chronic virus mimicry” to sustain inflammatory memory and ICB resistance. To test this, we genetically targeted RLRs, which are the pattern recognition receptors (PRRs) that likely recognize EREs to activate IFN signaling^38^. While we previously showed that RIG-I can contribute to ICB resistance^25^, MDA5 (*Ifih1*) was the only apparent RLR with significantly increased expression after chronic IFNG signaling **(Figure S2H)**. Deletion of MDA5 with or without RIG-I, but not other nucleic PRRs like PKR or STING, restored ICB response to ICB-relapsed Res 499 tumors **(Figure 2G; Figure S2I)**. Similarly, knockout of MDA5 in B16 melanoma or KP lung cancer cells that were rendered resistant by chronic IFNG stimulation also reversed resistance **(Figure 2H-I)**. Loss of resistance was additionally accompanied by loss of inflammatory memory. Here, RLR knockout in Res 499 cells abrogated persistent ISG-mem expression and the enhanced chromatin accessibility of its cis-REs at 72 hours after termination of IFNG stimulation **(Figure 2J-K)**. In contrast, ISG-mem and chromatin accessibility after 6 hours of IFNG stimulation was modestly impacted, while the kinetics of ISG-resolved was unaltered by RLR knockout. Unexpectedly, knockout of RLRs also abrogated the enhanced expression of EREs and dsRNAs associated with chronic IFNG stimulation and ICB resistance **(Figure 2L and 2B-C)**, suggesting the collapse of an entire feed-forward loop. In total, these findings suggest that resistance after ICB relapse or from chronic IFNG stimulation results in inflammatory memory through RLR signaling associated with de-repressed EREs and dsRNA. Specifically, an RLR-driven feed-forward loop maintains poised chromatin and expression of ISG-mem even after initial IFNG stimulation has terminated.

### Cancer cells with memory ISGs are found across human cancers and associate with CD8 T cell exhaustion

To understand the potential disease-relevance of ISG-mem, we assessed whether cancer cells expressing ISG-mem might exist in human cancers and if so, whether their presence correlates with features of immune dysfunction. For this, we utilized a comprehensive pan-cancer single-cell atlas^39^. Although cancer cells from tumors spanning different cancer types consistently showed low and homogenous expression of ISG-resolved and ISG-wave2, a discrete subpopulation demonstrated high ISG-mem expression **(Figure 3A-B; Figure S3A)**. To assess whether ISG-mem might be associated with immune dysfunction, we estimated tumor-infiltrating T cell subset proportions for all tumor samples containing both cancer cells and T cells. This revealed that in a pan-cancer analysis, the average cancer cell ISG-mem expression for individual tumors was strongly and positively correlated with the percentage of exhausted CD8 T cells (T_EX_) or progenitor T_EX_ (T_PEX_) but not most other T cell subtypes **(Figure 3C; Figure S3B)**. Although the sample size for many individual cancer types was low, the strength of this relationship between ISG-mem and T_EX_ was evident in specific cancer types such as kidney, breast, skin, and likely a subset of lung cancer **(Figure 3D)**. Thus, a minor subpopulation of cancer cells expressing ISG-mem are found across human cancers and correlate with features of CD8 T cell dysfunction.

**Figure 3.**
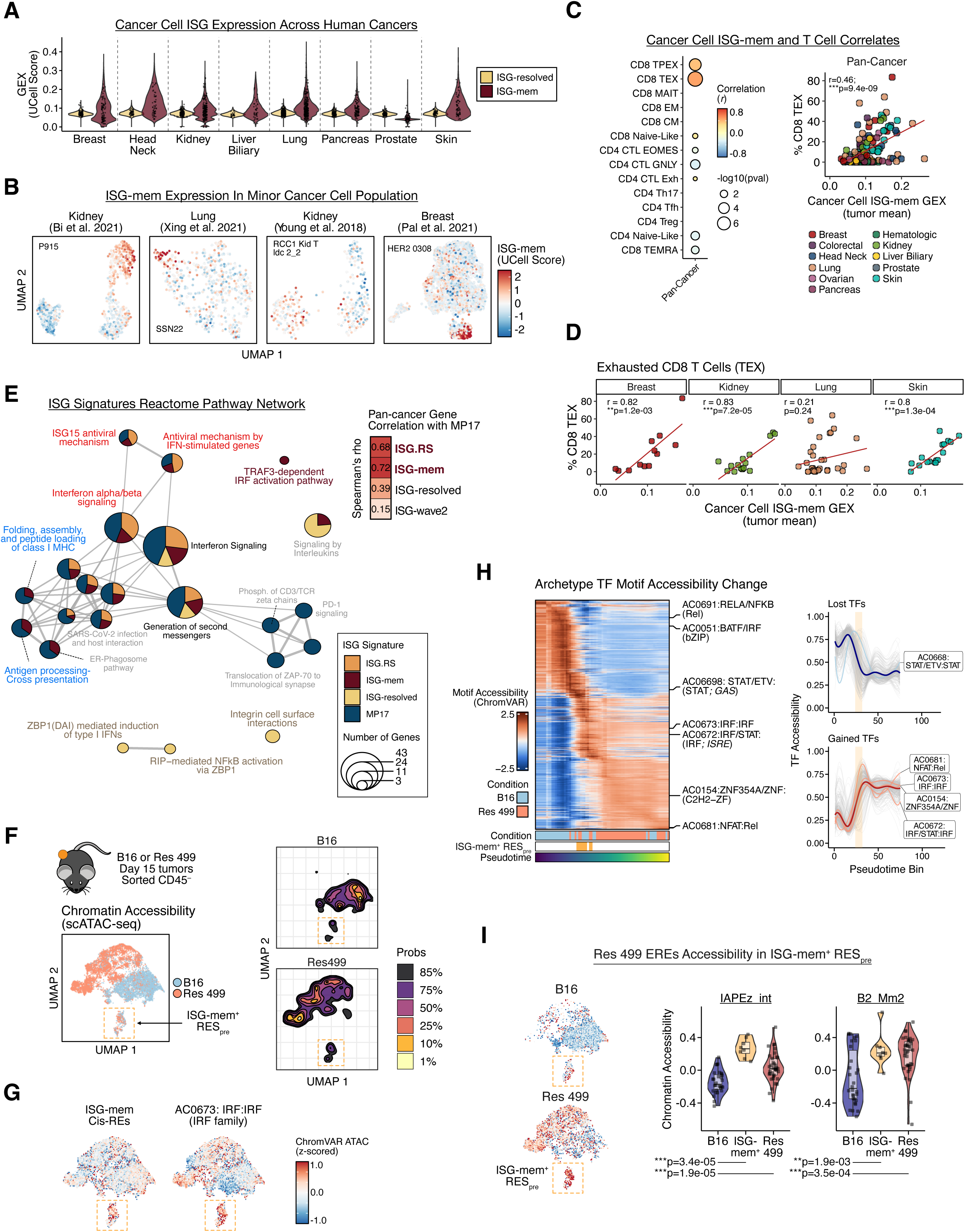
Subpopulation of cancer cells expressing memory ISGs are coupled to T cell exhaustion, found across human cancers, and exhibit resistance precursor properties. **A)** Expression of resolved and memory ISGs in cancer cells for the indicated human cancer types. **B)** Cancer cell memory ISG scaled expression on UMAP embeddings of representative human tumor samples. **C)** Pan-cancer summary of correlation between cancer cell memory ISG expression and abundance of the indicated T cell subtype (left) with Pearson correlation (color) and significance of the correlation coefficient (size) indicated. All tumor samples with >50 cancer cells and >50 T cells were used and shown in the scatter plot (right). **D)** Correlation between cancer cell memory ISG expression and proportion of exhausted CD8 T cells (TEX) is shown for the indicated example cancer types from the pan-cancer analysis from (C). **E)** Expression correlation and comparison of shared Reactome Pathways enriched in the indicated ISG signature. Pan-cancer gene expression correlations between indicated ISG signatures and the MP17 signature are shown in the heatmap along with rank correlation values (right inset). In the pathway network plot (left), all Reactome pathways significantly enriched in at least one ISG signature are plotted (pval < 0.05). Node colors indicate the ISG signatures significantly enriched for each pathway. Node size represents the number of ISG signature genes that belong to each pathway. Edge thickness represents gene similarity between pairs of Reactome pathways. **F)** UMAP embedding of chromatin accessibility features for B16 and Res 499 cells sorted from *in vivo* tumors, colored by condition (n=2 mice per condition; n=39,398 cells). Density contour plot representing regions of highest cell density from each condition is displayed (right). The ISG-mem^+^ RES_pre_ population is highlighted with an orange box. **G)** ChromVAR chromatin accessibility scores (bias-corrected deviation values) of the indicated chromatin signature overlaid on the UMAP embedding from (F). **H)** ChromVAR chromatin accessibility scores of Vierstra 2.0 archetype TF motifs in B16 and Res 499 cells ordered by pseudotime in a heatmap (left). Cells are binned and ChromVAR values are smoothed for visualization. TF motifs of interest are annotated. Co-regulated TF modules identified by hierarchical clustering (k=2) are summarized with a smoothed spline regression line with individual motif chromatin accessibility trends plotted in gray (right). Select TF motifs of interest are labeled. **I)** ChromVAR chromatin accessibility scores at Res 499-activated EREs overlaid on the UMAP embedding from (F). ISG-mem^+^ RES_pre_ population is highlighted by an orange box. Summary ATAC GSVA enrichment scores of chromatin accessibility at all expressed IAPEz-int and B2_Mm2 genome instances from pseudotime-binned *in vivo* cancer cells from the indicated condition (right). Unless otherwise indicated, P-values for two sample comparisons were determined by a two-sided t-test.

A consensus meta-program called MP17 was previously identified from the pan-cancer single-cell atlas as one of several highly prevalent gene programs accounting for tumor transcriptional heterogeneity in human cancers^39^. MP17 is characterized by coordinate upregulation of IFN response and MHC-II genes, and like ISG-mem, its expression arises from a subpopulation of cancer cells found across many common human cancers and is coupled to lower overall survival and therapy resistance^39^. In addition, we previously described an ISG signature that predicts clinical immunotherapy resistance after ICB^12^ and CAR-T cell therapy^13^ called the ISG.RS (ISG Resistance Signature) that can also be heterogeneously expressed in human tumors^25,40^. Thus, we sought to investigate the relationship between these different ISGs. Indeed, significant overlap exists between ISG-mem, MP17, and the ISG.RS **(Figure S3C)**, and the single-cell expression levels of these three signatures were strongly correlated across human cancers **(Figure 3E, top-right inset; Figure S3D)**. In contrast, despite also being ISGs, expression of ISG-resolved were weakly correlated with the other three ISG signatures. Pathway analysis revealed that ISG-mem, MP17, and ISG.RS share enrichment for IFN-I signaling and antiviral mechanisms **(Figure 3E, red labels)**, while ISG-mem and MP17 share antigen processing and presentation pathways (blue labels). However, only ISG-mem selectively enriches for IRF activation pathways (brown label), while ISG-resolved uniquely enriches for TNF/RIP-mediated NFKB signaling (beige labels). Thus, not only are ISG-mem enriched for immune evasion genes but are part of a specific and coordinated ISG program that is thematically-linked to antiviral IFN-I signaling and to immune dysfunction. Across human cancers, ISG-mem and these related ISGs are expressed by a small subpopulation of cancer cells and represent a major contributor to tumor heterogeneity.

### Epigenetically distinct cancer cells expressing memory ISGs resemble precursors to inflammatory memory and immunotherapy resistance

Like Res 499 tumors that relapsed after ICB, we recently reported that tumors from NSCLC patients that relapse after anti-PD1 acquire high levels of the same resistance-associated ISGs found in Res 499 tumors^9^. This raises the possibility that the pre-existing ISG-mem^+^ cells found in human cancers might give rise to ICB resistance. To investigate this notion, we examined whether a small population of ISG-mem^+^ cancer cells might pre-exist in mouse B16 tumors. Since the acquisition of ISG-mem is linked to epigenetic features of inflammatory memory, we generated single-cell chromatin accessibility (scATAC-seq) profiles of cancer cells from B16 and Res 499 tumors (n=39,398 cells from three independent experiments) **(Figure S3E)**. B16 and Res 499 epigenomes displayed extensive differences and separation into distinct clusters **(Figure 3F; Figure S3F)**. However, one small subpopulation of cells shared between B16 and Res 499 tumors exhibited both high chromatin accessibility scores at ISG-mem chromatin domains and IRF and STAT:IRF family motifs, but not at cis-REs for resolved ISGs or GAS motifs **(Figure 3G; Figure S3G)**. Thus, like with human tumors, a subpopulation of cancer cells expressing ISG-mem exist in heterogeneous tumors before immunotherapy treatment, raising the possibility that it may represent a putative ISG-mem^+^ resistance precursor (ISG-mem^+^ RES_pre_).

Since Res 499 relapsed from a B16 tumor after ICB, we investigated if pre-existing ISG-mem^+^ RES_pre_ cells might give rise to ICB-resistant cell states by first examining the inferred lineage relationship between B16 and Res 499 cancer cells. Pseudotime analysis rooted in cells with the lowest chromatin accessibility at Res 499 activated enhancers, which is a previous described set of cis-REs ascribed to Res 499 ICB resistance^25^, identified potential trajectories towards relapse^41^ (see Methods). Visualizing all archetype TF motifs along the relapse trajectory revealed a dramatically evolving epigenome characterized by two opposing TF regulatory programs **(Figure 3H)**. Most B16 cells shared the same chromatin state marked by high BATF/IRF (bZIP) motif activity. However, ISG-mem^+^ RES_pre_ cells resided at an inflection point in the predicted relapse continuum associated with major epigenetic changes, including the apparent onset of chronic IFNG-associated chromatin features prior to widespread increased accessibility of resistance-associated Res 499 activated enhancers **(Figure S3H)**. Like the majority of Res 499 cells, these ISG-mem^+^ RES_pre_ cells activated a suite of archetype TF motifs including those belonging to the IFN Regulatory Factor (IRF) family, containing IFN-sensitive Response Element (ISRE) motifs, or related to C2H2 zinc-finger TFs, which have been implicated in controlling ERE repression^42^ **(Figure S3H)**. Indeed, chromatin accessibility of EREs that were associated with chronic IFNG and resistant Res 499 tumors, such as IAPEz-int and B2 Mm2 families, was higher in the ISG-mem^+^ RES_pre_ population from Res 499 compared to B16 **(Figure 3I; Figure S3I)**. Thus, ISG-mem^+^ cells are found in mouse and human tumors and may represent precursor cells characterized by acquisition of elevated EREs, a subset of resistance-associated chromatin features, and the potential to transition into an ICB resistance state.

### Resistant tumors are epigenetically heterogenous with subclones differing in ICB response and features of inflammatory memory

To directly examine the developmental and resistance properties of ISG-mem^+^ RES_pre_ cells and other epigenetically-defined subpopulations from B16 and Res 499 tumors, we attempted to isolate these subpopulations by generating single-cell derived populations (SCPs) from B16 and Res 499 cells **(Figure 4A)**. Using chromatin accessibility profiles from *in vivo* SCP-derived tumors, we mapped each SCP back to single-cell chromatin state(s) of B16 and Res 499 tumors. Assessment for phenotypic and molecular heterogeneity revealed that all B16-derived SCPs were equally or more sensitive to ICB compared to parental B16 **(Figure S4A-B)** and exhibited little or indistinct epigenetic heterogeneity **(Figure S4C)**. In contrast, four out of fifteen Res 499 SCPs were sensitive to ICB **(Figure S4D, blue and orange)**, while the rest were resistant **(Figure S4D, red and pink)**. Accordingly, the epigenomes of sensitive Res 499 SCPs resembled bulk B16 parental tumors, while resistant Res 499 SCPs were epigenetically more similar to Res 499 bulk tumors **(Figure 4B-C)**. Using hierarchical clustering and principal component analysis of chromatin accessibility profiles, we identified five chromatin states among the Res 499 SCPs: two states corresponding to resistant SCPs (Epi-Res1, Epi-Res2), two states associated with sensitive SCPs (Epi-Sen1 and Epi-Sen2, or collectively Epi-Sen1/2) that resembled the B16-like sensitive state, and one unique state belonging to Res 499 SCP 11 (Epi-SCP11). Mapping the SCP chromatin states to single-cell chromatin accessibility features of B16 and Res 499 cells sorted from *in vivo* tumors revealed that these five states accounted for much of the epigenetic variation in Res 499 tumors **(Figure 4D-E; Figure S4E-F)**.

**Figure 4.**
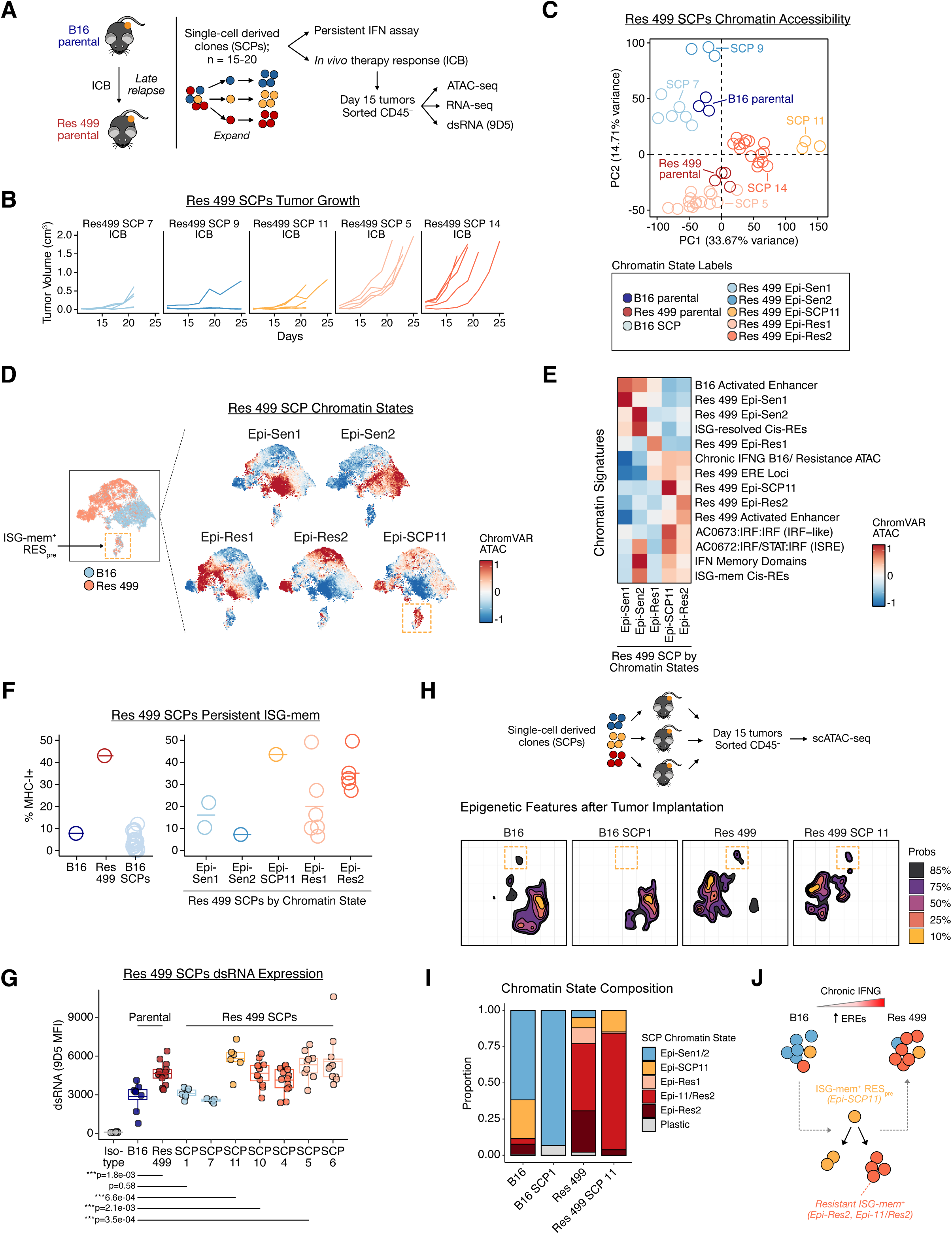
Subpopulations of epigenetically distinct cancer cells display a spectrum of phenotypic features coupled to ICB sensitivity and resistance. **A)** Experimental design for phenotypic and multi-omic profiling of single-cell derived populations (SCPs) from B16 and Res 499 parental tumors. **B)** Tumor growth of mice implanted with representative Res 499 SCPs and treated with anti-CTLA4 + anti-PDL1 (ICB) (n=5-10 per condition). See Figure S4D for tumor volumes of all SCPs. **C)** Principal component analysis of genome-wide chromatin accessibility profiles from Res 499 SCPs. SCPs are color-coded by chromatin state that corresponds to ICB response phenotype. **D)** ChromVAR chromatin accessibility scores of Res 499 SCP chromatin state signatures overlaid on the UMAP embedding from Figure 3F. **E)** Heatmap of average ChromVAR chromatin accessibility scores from Res 499 SCPs grouped by the indicated chromatin signatures. **F)** Percent MHC-I-positive cancer cells at 72 hours washout following IFNG stimulation for the *in vitro* persistent ISG-mem assay described in Figure 1J. Res 499 SCPs are grouped by chromatin state. **G)** MFI of dsRNA staining using the 9D5 antibody from cancer cells sorted from the indicated tumors (n=10 mice per condition). **H)** Density contour plot of UMAP embedded chromatin accessibility features from the indicated B16, Res 499, or SCP tumor. Location of the ISG-mem^+^ RES_pre_ population is highlighted by the orange box. **I)** Proportion of cancer cells in each tumor from (H) assigned to the indicated SCP chromatin state annotation. **J)** Schema depicting the lineage properties of ISG-mem^+^ RES_pre_ cells in B16 and Res 499 tumors. Unless otherwise indicated, P-values for two sample comparisons were determined by a two-sided t-test.

To determine whether differences in ISG-mem epigenetic features were functionally linked to inflammatory memory, we tested each SCP for persistent ISG-mem expression after termination of IFNG signaling. Indeed, Epi-Res2 SCPs, but not B16-like SCPs, maintain persistent MHC-I expression 72 hours after IFNG stimulation, a surrogate for acquisition of ISG-mem **(Figure 4F)**. Epi-Res2 SCPs also exhibited elevated EREs and dsRNA, consistent with chronic virus mimicry promoting inflammatory memory in these SCPs **(Figure 4G; Figure S4G-H)**. Consistent with these features, Res 499 SCPs with the Epi-Res2 state also displayed high chromatin accessibility for ISG-mem cis-REs and Res 499 EREs, in addition to cis-REs acquired after chronic IFNG stimulation, Res 499 activated enhancers, and IRF and ISRE motifs **(Figure 4E; Figure S4F)**. The Epi-Res1 SCPs lacked elevated accessibility for many of these epigenetic features and had more modest persistence of MHC-I, suggesting that it captures an alternative and possibly IFN-independent resistance mechanism. Thus, Res 499 cells possessing a stable Epi-Res2 chromatin state may account for ICB resistance resulting from chronic IFNG stimulation promoting inflammatory memory and acquisition of ISG-mem expression.

The Epi-SCP11 chromatin state represents Res 499 SCP 11, a rare SCP that has a unique molecular and phenotypic profile **(Figure 4C)**. Res 499 SCP 11 displays many of the same collection of features as cells with the Epi-Res2 resistant state – namely, chronic virus mimicry and persistent ISG-mem expression **(Figure 4F-G; Figure S4G-H)**. However, Res 499 SCP 11 shows incomplete resistance-associated epigenetic features by lacking, on average, some Res 499 activated enhancers that are linked to the Epi-Res2 chromatin state **(Figure 4E; Figure S4I)**. Accordingly, Res 499 SCP 11 tumors were more sensitive to ICB than tumors from Epi-Res2 SCPs **(Figure 4B)**. This suggests that cells with an Epi-SCP11 chromatin state may belong to the ISG-mem^+^ RES_pre_ putative precursor population that gives rise to resistant chromatin states. Indeed, the strongest enrichment of the Epi-SCP11 chromatin state was observed in the ISG-mem^+^ RES_pre_ population that was shared between B16 and Res 499 **(Figure 4D)**. Other SCP chromatin states enriched within distinct and mostly non-overlapping epigenetic populations but not within the ISG-mem^+^ RES_pre_ population. Notably, cells with other SCP chromatin states, especially Epi-Res2, also demonstrated variable enrichment for the Epi-SCP11 chromatin state, suggesting retained accessibility possibly from a developmental relationship. Consistent with this, implantation of Res 499 SCP 11 cells into mice predominantly reconstituted populations either enriched in the Epi-Res2 chromatin states or cells enriched in both Epi-SCP11 and Epi-Res2 chromatin states **(Figure 4H-I)**. Cells with B16-like Epi-Sen1/2 chromatin states rarely emerged from Res 499 SCP 11 *in vivo*, even though a small population with a B16-like chromatin state were found in Res 499 tumors. In contrast, implantation of B16 SCP 1 into mice gave rise almost exclusively to populations with the B16-like Epi-Sen1/2 chromatin states but not Epi-SCP11 or any other Res 499 chromatin states, even though B16 tumors were comprised of minor populations of cells with Epi-SCP11 and Res 499 chromatin states. Thus, lineage-restricted SCP chromatin states appear to exist in B16 and Res 499 tumors. In total, these data suggest that Res 499 SCP 11 is from the ISG-mem^+^ RES_pre_ population shared between B16 and Res 499 tumors and functions as a precursor for ICB resistant states, possibly driven by chronic IFNG signaling and chronic virus mimicry **(Figure 4J)**.

### Chronic antiviral signaling through an MDA5 feedforward loop is required for relapse of precursors expressing memory ISGs

Our results demonstrate that chronic IFNG and RLR signaling in Res 499 cells, possibly involving elevated EREs and dsRNAs, are responsible for inflammatory memory and ICB resistance. Notably, EREs appear more elevated in ISG-mem^+^ RES_pre_ cells from Res 499 compared to B16 tumors **(Fig. 3I)**, suggesting chronic IFNG and RLR signaling might drive relapse of ISG-mem^+^ RES_pre_ cells. To examine this, we first chronically stimulated Res 499 SCP 11 cells with IFNG *in vitro* followed by tumor implantation into mice, and for comparison, repeated this with the sensitive Res 499 SCP 9. For Res 499 SCP 11, chronic IFNG resulted in accelerated development of ICB resistance as measured by tumor growth and survival, while no discernible effects were apparent for Res 499 SCP 9 **(Figure 5A-B)**. Conversely, we examined the effect of MDA5 deletion from Res 499 SCP 11 cells prior to tumor implantation into mice. While Res 499 SCP 11 wildtype tumors were initially sensitive to ICB but then began relapsing at around day 25, knockout of MDA5 prevented relapse for up to 40 days **(Figure 5C)**. Thus, these data suggest that relapse of ISG-mem^+^ RES_pre_ cells is controlled by chronic IFNG and RLR signaling.

**Figure 5.**
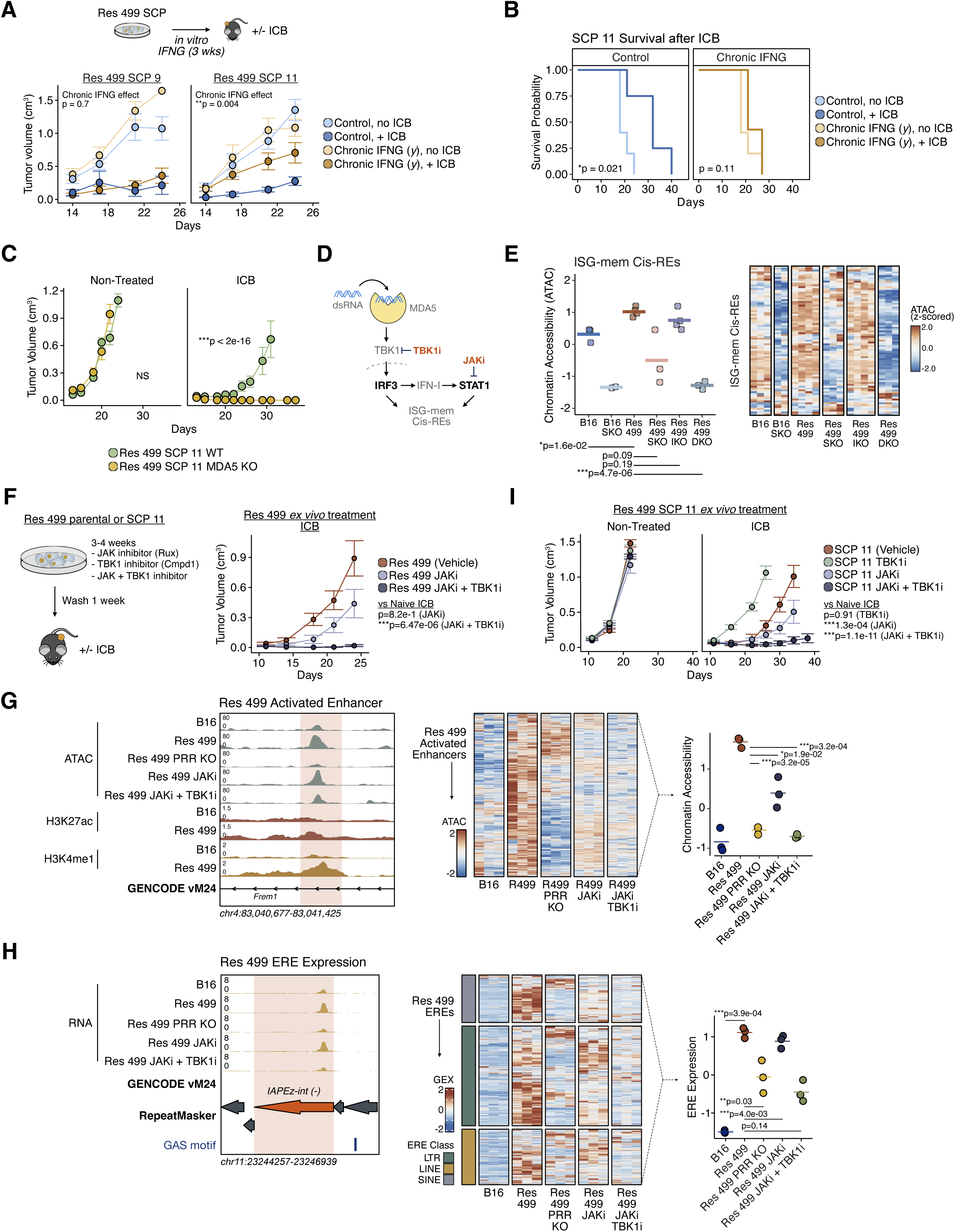
Precursor cells expressing memory ISGs acquire resistance through an MDA5-mediated feedforward mechanism abrogated by JAK plus TBK1 inhibitors. **A)** Tumor growth after implantation of ISG-mem^+^ Res 499 SCP 11 cells either control-treated or treated with chronic IFNG *in vitro* for 3 weeks. For comparison, Res 499 SCP 9 cells enriched for the Epi-Sen2 chromatin state were also used. Mice were either untreated (n=10 per condition) or treated with anti-PDL1 + anti-CTLA4 (ICB) (n=19-37 per condition). P-value indicates significance for an effect of chronic IFNG on modifying ICB response. **B)** Survival of mice from (A) implanted with Res 449 SCP 11 cells either control-treated or treated with chronic IFNG for 3 weeks *in vitro*. **C)** Tumor growth from mice implanted with Res 499 SCP 11 wildtype (WT) or MDA5 knockout (KO) cells either non-treated (n=5 per condition) or treated with ICB (n=10 mice per condition). **D)** Schema of MDA5 signaling pathway and effect of JAK inhibitor (JAKi) or TBK1 inhibitor (TBK1i). **E)** Summary ATAC GSVA enrichment score (left) and heatmap (right) of ISG-mem Cis-REs from B16 or Res 499 cancer cells with or without STAT1 KO (SKO), IRF3 KO (IKO), or STAT1 plus IRF3 KO (DKO) sorted from *in vivo* tumors (n=2-4 mice per condition). Individual biological replicates are shown in the heatmap. **F)** Experimental design for JAKi + TBK1i treatment of Res 499 or Res 499 SCP 11 cells (left). Tumor growth from mice implanted with Res 499 cells after *in vitro* treatment with JAKi, TBK1i, or JAKi + TBK1i. Mice were treated with or without ICB (n=8 mice per condition). **G)** Heatmap and summary plot of chromatin accessibility scores for Res 499 activated enhancers (right) and representative chromatin accessibility (ATAC), H3K27ac and H3K4me1 CUT&RUN tracks (left) from B16 or Res 499 cancer cells with or without MDA5/RIG-I KO (PRR KO) or expanded *in vitro* with or without JAKi +/- TBK1i, sorted from *in vivo* tumors. Heatmap shows individual biological replicates. **H)** Same as (G) but for RNA expression of EREs upregulated in Res 499 tumors. GENCODE vM24 gene and RepeatMasker annotations, and instances of the GAS TF archetype motif (AC0668) are indicated. Individual biological replicates are shown in the heatmap. **I)** Tumor growth from mice implanted with Res 499 SCP 11 cells after *in vitro* treatment with JAKi, TBK1i, or JAKi + TBK1i. Mice were treated with or without ICB (n=8 mice per condition). Unless otherwise indicated, P-values for two sample comparisons were determined by a two-sided t-test. P-values for tumor growth were determined by a mixed-effect regression model, and for survival by a log-rank test. Error bars represent standard errors.

ISG-mem^+^ RES_pre_ cells lie at an inflection point between B16 and Res 499 that is characterized by major epigenetic changes associated with IRF and STAT:IRF family TF motifs **(Figure 3H)**. RLRs activate the serine/threonine kinase TBK1 to phosphorylate IRF3, resulting in IFN-I production that further signals through JAK kinases to activate STAT1^38^. Hence, these observations suggest that RLR-dependent relapse and the development of epigenetic changes in ISG-mem^+^ RES_pre_ cells might be driven by STAT1 and IRF3. Moreover, the combination of JAK and TBK1 inhibition could be a pharmacological approach to recapitulate effects of RLR deletion **(Figure 5D)**. Indeed, simultaneous deletion of both STAT1 and IRF3, but not deletion of each individually, near-completely reversed the enhanced chromatin accessibility at ISG-mem cis-REs similar to STAT1 knockout in B16 tumors that do not exhibit inflammatory memory **(Figure 5E)**. STAT1 and IRF3 deletion also abrogated dsRNA levels to B16 tumor levels **(Figure S5A)**. In contrast, no impact on cis-REs for ISG-resolved was observed **(Figure S5B)**. To examine the combination of JAK and TBK1 inhibitors (JAKi + TBK1i), we expanded Res 499 cells for 3 weeks *in vitro* in the presence of no drug (vehicle), a JAK inhibitor (ruxolitinib), and/or a TBK1 inhibitor (Cmpd1), and then washed for 1 week prior to *in vivo* tumor implantation. Treatment with the JAKi + TBK1i potently resensitized Res 499 tumors to ICB **(Figure 5F)** and largely reversed the enhanced chromatin accessibility of Res 499 activated enhancers marked by H3K4me1, including those found in ISG-mem^+^ RES_pre_ (Res 499 SCP 11) cells, to levels similar to B16 or Res 499 with RLR knockout **(Figure 5G; Figure S5C-D)**. This was accompanied by a similar abrogation of elevated EREs **(Figure 5H)**. Notably, JAKi + TBK1i was more effective than JAK inhibitor alone for all of these effects, which is consistent with how STAT1 plus IRF3 knockout was more effective than STAT1 knockout alone at reversing enhanced chromatin accessibility of resistant Res 499 tumors. We next repeated the *in vitro* treatment using JAKi + TBK1i for Res 499 SCP 11 cells prior to tumor implantation into mice. Like with MDA5 deletion, this combination effectively prevented ICB relapse, while JAKi alone or TBK1i alone led to more modest improvement or even worse response, respectively **(Figure 5I)**.

In total, these findings suggest that relapse of ISG-mem^+^ RES_pre_ cells is controlled by chronic IFNG and MDA5 signaling that results in STAT1 and IRF3 driving epigenetic changes of inflammatory memory. Pharmacological inhibition of JAK and TBK1 collapses the MDA5-mediated feed-forward loop that sustains inflammatory memory, highlighting a therapeutic approach to prevent relapse of the ISG-mem^+^ RES_pre_ population.

### JAK and TBK1 inhibition prevents development of resistant epigenetic states

Inflammatory memory can be inherited across cell divisions and promote a differentiation bias in hematopoietic stem and progenitor cells^43,44^. Therefore, we wondered whether JAK plus TBK1 inhibition prevents relapse by interfering with inflammatory memory in ISG-mem^+^ RES_pre_ cells, hence perturbing the inheritance of resistance-related epigenetic features. To assess this, we utilized cell barcoding to enable simultaneous lineage tracing and profiling of epigenetic states^45^. Bulk Res 499 cells were transduced with the Watermelon barcode library and treated with either vehicle or JAKi + TBK1i for 3 weeks *in vitro*. Following a one-week washout, cells were collected for scATAC-seq and lineage barcode read-out **(Figure 6A, Figure S6A-C)**, allowing us to capture approximately 1500 clones *in vitro*. While most chromatin signatures exhibited stochastic chromatin accessibility within lineage barcodes (clones), chronic IFNG signatures, SCP chromatin signatures, IFN memory domains, and ISRE chromatin accessibility were more similar within clones than across clones, confirming these epigenetic features are stably heritable over cell divisions *in vitro* and/or *in vivo* **(Figure 6B, Figure S6D)**. Next, we annotated each cell by SCP chromatin state (see Methods). This demonstrated that most cells mapped to one chromatin state, some mapped to both Epi-SCP11 and Epi-Res2 chromatin states (Epi-11/Res2), while some did not clearly map to a single state (Plastic). Examination of the cell composition of lineage barcodes revealed that most lineage barcodes were comprised of multiple SCP chromatin states and nearly all contained cells with the Epi-SCP11 chromatin state, consistent with cells in each clone potentially originating from ISG-mem^+^ RES_pre_ cells **(Figure 6C)**. In the vehicle condition, most cells within a lineage barcode possessed either resistant Epi-Res2 or the sensitive Epi-Sen1/2 chromatin state. In the JAKi + TBK1i condition, examination of the same clones revealed a dramatic shift in the *in vitro* clonal composition, resulting in a reduction in the proportion of cells with an Epi-Res2 or Epi-11/Res2 chromatin state and a concomitant increase in cells with an Epi-Sen1/2 chromatin state **(Figure 6C-D)**. This altered clonal composition was not explained by increased cell death, as a strong correlation in clone size between vehicle and JAKi + TBK1i treated cells was observed *in vitro* regardless of its dominant SCP chromatin state composition **(Figure 6E, Figure S6E)**. However, JAKi + TBK1i noticeably diminished ISG-mem chromatin accessibility and reciprocally increased ISG-resolved chromatin accessibility in ISG-mem^+^ RES_pre_ cells **(Figure 6F)**. Thus, ISG-mem and SCP chromatin states are clonally heritable, and JAKi + TBK1i treatment appears to perturb both chromatin accessibility of ISG-mem in ISG-mem^+^ RES_pre_ cells and possibly lineage relationships between these precursors and cells with sensitive and resistant epigenetic states.

**Figure 6.**
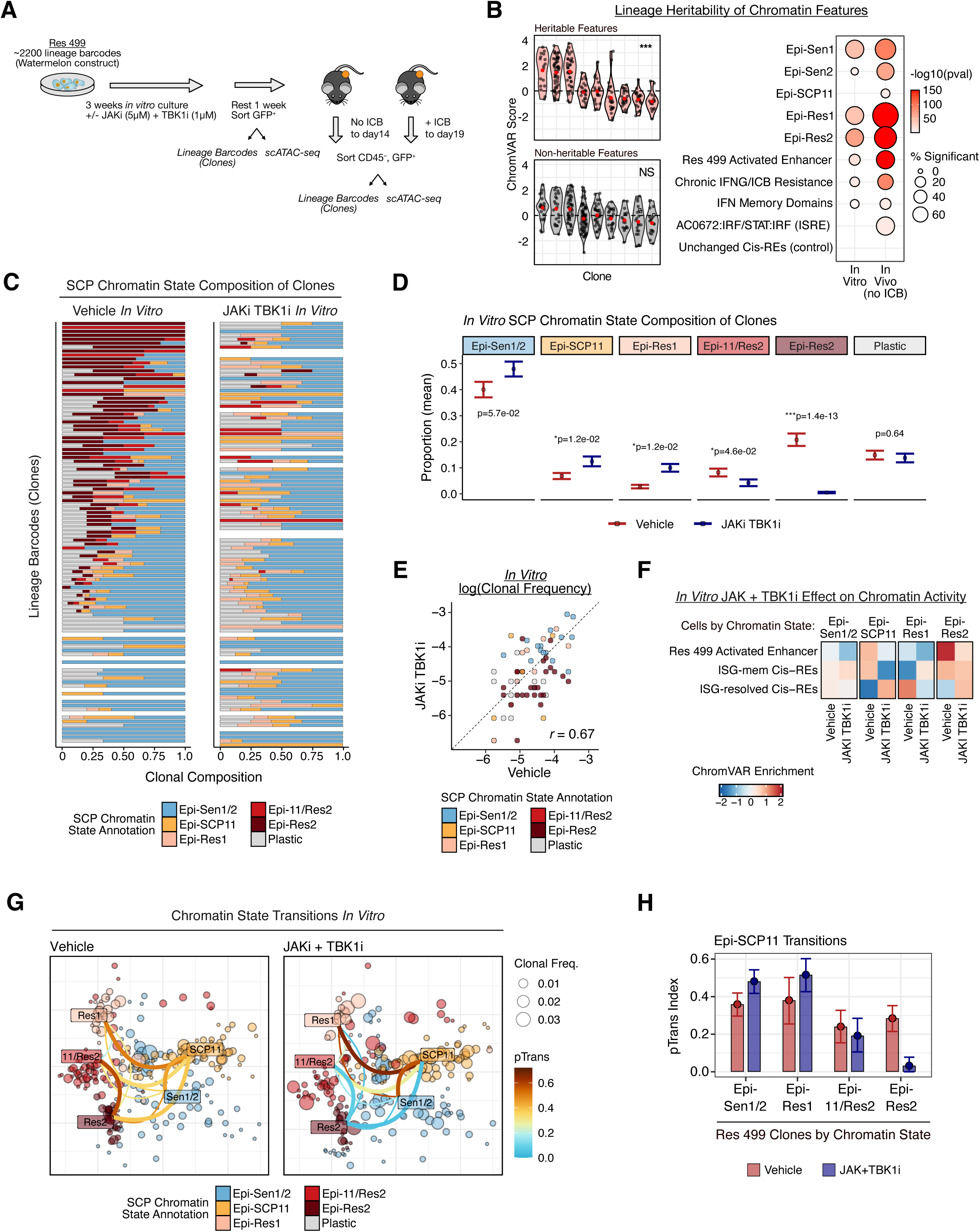
JAKi plus TBK1i alters differentiation dynamics of memory ISG-expressing precursors to prevent development of resistant epigenetic states. **A)** Experimental design to analyze clonal dynamics, lineage properties, and *in vivo* ICB tumor response of Res 499 cells treated with or without JAKi + TBK1i *in vitro*. **B)** Example plots of a heritable (left, top) and non-heritable (left, bottom) chromatin signatures where y-axis is chromatin accessibility score and x-axis is cells grouped by barcode. Dot plot of significance values from one-way ANOVA testing for clonal heritability of indicated chromatin signatures (right). Color of dots represents P-value and size represents percent of clones showing significant heritability of chromatin signatures. **C)** SCP chromatin state composition for Res 499 clones (i.e., cells with same lineage barcode). Shown are the proportion of Res 499 cells within each lineage barcode (rows) with the indicated SCP chromatin state (color-coded) after *in vitro* treatment with vehicle or JAKi + TBK1i. The top lineage barcodes by frequency are shown. **D)** Average proportion of cells with the indicated SCP chromatin states within individual lineage barcode averaged across all Res 499 clones. **E)** Lineage barcode frequency for Res 499 cells treated *in vitro* with vehicle compared to frequency after JAKi + TBK1i. Each clone is colored by its dominant SCP chromatin state annotation. **F)** Mean ChromVAR chromatin accessibility scores (bias-corrected deviation scores) for the indicated chromatin signatures in cells annotated by SCP chromatin state and grouped by treatment condition. **G)** Lineage barcode sharing between clones annotated by SCP chromatin states are overlaid on a UMAP embedding for chromatin accessibility features. Clones from each treatment condition are separately shown. Edges connecting clones grouped by their dominant SCP chromatin state (nodes) are colored by a pairwise transition-index (pTrans-index), with higher scores indicating greater barcode sharing between groups (a measurement of developmental relatedness). Edges involving Epi-SCP11 or between Epi-11/Res2 and Epi-Res2 groups are thicker for easier visualization. Each lineage barcode is plotted as the average of their UMAP dimensions and colored by its dominant SCP annotation. Dot size indicates lineage barcode frequency. **H)** Bootstrapped pTrans-index values between clones with a dominant Epi-SCP11 chromatin state and clones with the other dominant SCP chromatin states. Results are color-coded by treatment condition. Error bars represent the 95% confidence interval. P-values for two sample comparisons were determined by a two-sided t-test.

To quantitate changes in developmental relatedness of ISG-mem^+^ RES_pre_ cells with cells possessing other epigenetically distinct states, we used a pairwise transition index (pTrans-index) to assess the degree of barcode sharing^46^. In the vehicle condition, strong developmental relatedness was noted between cells with the Epi-SCP11 chromatin state (ISG-mem^+^ RES_pre_ cells) and cells with the other major SCP chromatin states – Epi-Res2, Epi-11/Res2, Epi-Res1, and Epi-Sen1/2 **(Figure 6G, Figure S6F)**. However, JAKi + TBK1i treatment led to greatly increased barcode sharing between ISG-mem^+^ RES_pre_ cells and Epi-Sen1/2 cells and to reduced sharing with Epi-Res2 cells **(Figure 6G-H)**. Sharing between ISG-mem^+^ RES_pre_ cells and Epi-11/Res2 was more modestly decreased, while sharing with Epi-Res1 cells increased with JAKi + TBK1i, likely due to continued differentiation toward this alternative and inflammatory memory-independent resistant state. Together, these data suggest that the clonal composition differences and ICB sensitivity observed with *in vitro* JAK plus TBK1 inhibition can be explained by perturbed differentiation dynamics of ISG-mem^+^ RES_pre_ cells. Specifically, JAK plus TBK1 inhibition decreases ISG-mem chromatin accessibility in ISG-mem^+^ RES_pre_ cells, blocking their differentiation toward resistant Epi-Res2 epigenetic states, which directly pivots or indirectly results in development toward sensitive states.

We next determined the consequences of perturbing clonal differentiation dynamics after JAKi + TBK1i on tumor heterogeneity and ICB response *in vivo*. After implantation into mice, vehicle-treated Res 499 cells gave rise to tumors that were expectedly resistant to ICB **(Figure 7A)**. Examination of *in vivo* clonal composition revealed an overall decrease in the proportion of Epi-Sen1/2 chromatin states and an increase in Epi-11/Res2 and/or Epi-Res2 states when compared to the *in vitro* clonal composition prior to implantation, an effect that was further exaggerated by ICB **(Figure 7B, left; Figure 7C and 6D)**. In contrast, implantation of JAKi + TBK1i treated cells into mice resulted in a large reduction in the clonal composition of cells with Epi-11/Res2 chromatin state, compared to vehicle-treated cells, despite exposure to JAKi + TBK1i over 3 weeks prior **(Figure 7C)**. After ICB therapy, JAKi + TBK1i led to a reduction in the clonal composition of both Epi-11/Res2 and Epi-Res2 chromatin states, and many clones became undetectable **(Figure 7B, right; Figure S6E)**, consistent with JAKi + TBK1i preventing tumor relapse **(Figure 7A)**. To gain additional insight into the nature of the clones that disappeared or decreased in frequency, we assigned clones a dominant SCP chromatin state based on their *in vitro* status prior to JAKi + TBK1i treatment **(Figure S7A)**. Tracking the fate of these clones *in vivo* after tumor implantation revealed that Epi-Sen1/2 chromatin state clones diminished in frequency after ICB therapy, while clones initially assigned to the Epi-Res2 state maintained their abundance under ICB pressure **(Figure 7D)**. However, with JAKi + TBK1i treatment prior to implantation, ICB effectively depleted these Epi-Res2 clones. Examination of JAKi + TBK1i treated cells that persisted in the tumor after ICB revealed that a large proportion possessed the resistant Epi-Res1 chromatin state **(Figure 7E; Figure S7B-C)** that is unrelated to inflammatory memory **(Figure 4E)**, while vehicle-treated cells that persisted in the tumor were predominantly Epi-Res2 cells. In total, these data suggest that JAK plus TBK1 inhibition directly blocks ISG-mem^+^ RES_pre_ cells from developing into ISG-mem^+^ resistant epigenetic states, diminishing the likelihood of tumor relapse after ICB **(Figure 7F)**.

**Figure 7.**
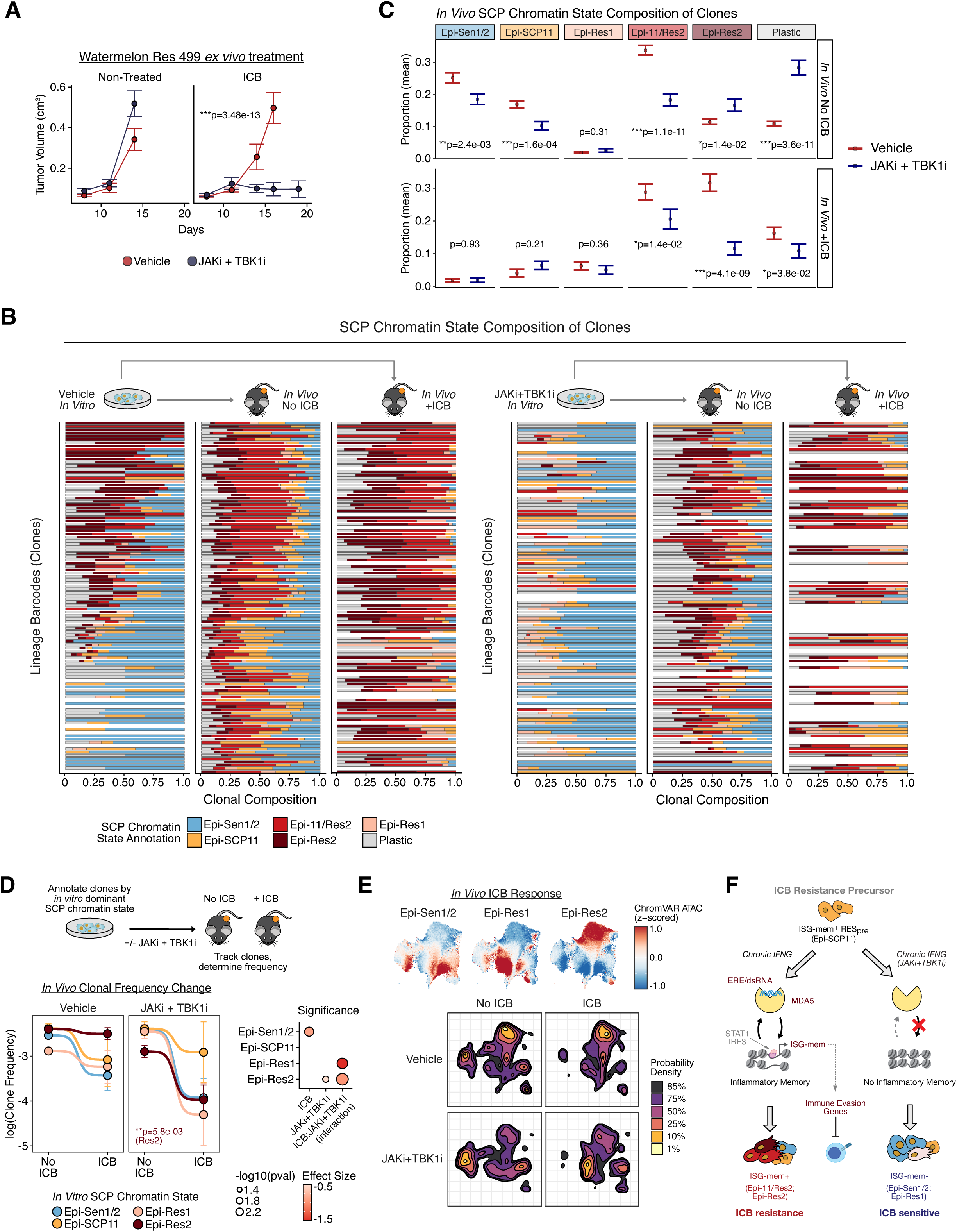
JAKi plus TBK1i restores ICB response of tumors that have acquired resistance through chronic IFNG and inflammatory memory. **A)** Tumor growth from mice implanted with Watermelon-barcoded Res 499 cells expanded *in vitro* with no treatment (Vehicle) or in the presence of JAKi + TBK1i for 3 weeks. Mice were left untreated (n=5 per condition) or treated with aCTLA4 + aPDL1 (n=10 mice per condition). **B)** Change in SCP chromatin state composition of Res 499 clones (rows) treated with vehicle (left) or JAKi + TBK1i (right) and then implanted into mice treated with or without ICB. Lineage barcodes with highest frequencies were considered. The *in vitro* data plots shown are the same as Fig. 6C. **C)** SCP chromatin state composition of clones for each indicated treatment condition averaged across all lineage barcodes displayed in (B). **D)** Impact of ICB on persistence of Res 499 clones annotated by initial dominant SCP chromatin state. Clones were annotated based on dominant SCP chromatin state at starting *in vitro* time point (Fig. 6A) and then expanded with (right) or without (left) JAKi + TBK1i. These clones were then tracked *in vivo* after tumor implantation into mice treated with and without ICB. The significance of the effect of ICB, JAKi + TBK1i, or the interaction between ICB and JAKi + TBK1i on *in vivo* clonal frequencies are shown in the dot plot (right) with size indicating P-value and color representing effect size. **E)** ChromVAR chromatin accessibility scores for SCP chromatin states in Res 499 cells overlaid on a UMAP embedding of chromatin accessibility features (top). Lineage barcoded cells were first expanded with or without JAKi + TBK1i *in vitro* prior to tumor implantation into mice that were then treated with or without ICB. Density plot of the UMAP embedding for each *in vivo* condition demonstrates the distribution of persistent clones in the tumor. **F)** Model for clonal differentiation dynamics and ICB resistance driven by inflammatory memory. P-values for two sample comparisons were determined by a two-sided t-test. P-values for tumor growth were determined by a mixed-effect regression model. Error bars represent standard errors.

## DISCUSSION

With the success of cancer immunotherapies such as immune checkpoint blockade, acquired resistance has become an emerging obstacle to durable response and hence long-term overall survival^8^. Our study suggests that an important mechanism of acquired resistance may originate from a pre-existing subpopulation of cancer cells that evolve into resistant states. This subclonal evolution can be driven by chronic inflammatory signaling, particularly IFNG, that leads to the development of inflammatory memory. In normal tissues such as epithelial and hematopoietic compartments, inflammatory memory typically involves stem and progenitor cells, and can represent an adaptive response to chronic infection and chronic disease^23,24^. Inflammatory memory may bias the differentiation of these normal tissue stem and progenitor cells to favor cell lineages that help to restore homeostasis, while also augmenting expression of immune modulatory genes to mitigate detrimental effects that inflammation has on tissue homeostasis. Our study demonstrates that subpopulations of cancer cells with precursor properties may similarly develop inflammatory memory for purposes of biasing differentiation toward therapy resistant states and to enhance expression of genes needed to avoid immune-mediated attack. The development of inflammatory memory by cancer cells is associated with the acquisition of poised chromatin and enhanced transcriptional response for a subset of memory ISGs that can persist even after initial IFNG stimulation is terminated. Thus, the acquisition of an epigenetically-encoded recall response to prior chronic IFNG stimulation by a small subpopulation of cancer cells may prime tumors for developing ICB resistance. The pervasive existence of a minor ISG-mem^+^ subpopulation across human cancers suggest that these cells may constitute a clinically relevant nidus for acquired resistance. Targeting a core pathway involved in sustaining inflammatory memory using JAKi + TBK1i prevents ISG-mem^+^ cells from giving rise to resistant epigenetic states, highlighting a pharmacological approach to potentially promote durable ICB efficacy.

Inflammatory memory, whether in normal tissues or cancer cells, requires a mechanism to sustain long-lasting changes to cells even after initial inflammatory stimuli has ceased^23^. A positive feedback and/or feed-forward biological circuit would serve such a purpose; however, a molecular basis for inflammatory memory has been unclear. Several studies, including our previous work, have suggested that inflammatory memory may involve nucleic acid pattern recognition receptors^25,47^ and/or STAT-family transcription factors^23,25,47,48^. Our current findings extend these observations and shed insight into potential mechanisms. We show a strong coupling between epigenetic, transcriptional, and functional features of inflammatory memory with the transcriptional de-repression of EREs and an accumulation of cytoplasmic dsRNA. Such dsRNA, which may represent ERVs or other transcripts with inverted SINE repeats^38^, are thought to bind to RNA-sensing PRRs like MDA5 that directly activates IRF3 through TBK1 and indirectly activates STAT1 through autocrine IFN-I and JAK kinases. Our findings that MDA5 deletion, STAT1 and IRF3 co-deletion, and JAKi + TBK1i all similarly abrogate epigenetic, transcriptional, and functional features of inflammatory memory reveal the nature of a positive feedback or feed-forward mechanism for ICB resistance – chronic virus mimicry through EREs and RNA-sensing PRRs. How chronic IFNG signaling initiates this memory-promoting mechanism is currently unclear. Some EREs may be ISGs themselves, or certain ISG transcripts may have a predilection for forming dsRNA^38^. For example, cis-REs for EREs can have an enrichment in TFs for STAT1, and introns can contain inverted SINE repeats that promote dsRNA formation^49,50^. In cancer, RNAs called SPARCS with ERE cis-REs in the 3’ UTR can initiate antisense strand transcription, resulting in dsRNA formation^51^. Regardless of the mechanism for how dsRNAs are formed, epigenetic remodeling leading to transcriptional de-repression of dsRNA-forming transcripts likely play an important role as well, as EREs are typically silenced through H3K9me3 and other silencing mechanisms^52^. Our findings that STAT1 and IRF3 together maintain chromatin accessibility not only of inflammatory memory domains but also of chromatin regions surrounding EREs may unveil why MDA5 signaling is an important component of the positive feedback loop – MDA5 controls chromatin accessibility and transcriptional activation of ligands that stimulate it.

In ICB-resistant cancer cells, the transcriptional and chromatin features of inflammatory memory are accompanied by functional properties that include persistent expression of memory ISGs even after the termination of IFNG stimulation. This persistence of transcription is likely due to autocrine IFN-I production that we previously showed is significantly elevated after ICB relapse due at least in part to OAS1^25^, another dsRNA PRR and memory ISG. Memory ISGs are also significantly enriched in bone fide immunotherapy resistance genes that were identified by *in vivo* genome-wide CRISPR screening. This suggests that one important function of memory ISGs is to promote immune evasion of ISG-mem^+^ RES_pre_ cells, as well as its progeny with the Epi-Res2 chromatin state, which likely captures cis-REs for additional immune evasion genes. It is notable that cancer cells with sensitive SCP chromatin states also have high chromatin accessibility for ISG-mem cis-REs; however, unlike resistant cells with Epi-Res2, these sensitive cell states additionally have high chromatin accessibility for ISG-resolved cis-REs. This suggests that ICB resistance may require a downregulation of ISG-resolved in addition to enhanced expression of ISG-mem. One possibility is that resistance entails not only augmented and persistent expression of immunosuppressive ISG-mem but also a “culling” of ISGs like ISG-resolved, which might be collectively more immunostimulatory due to genes in the TNF/RIP-mediated NFkB pathway, for example. In fact, negative feedback mechanisms to suppress aspects of IFN signaling are well known and can be controlled by negative regulators such as *Usp18*^33^, which is also an ISG-mem gene. Opposing immune effects of different ISG subsets not only could contribute to how IFNs can have opposing roles in immune regulation but also why different subsets of ISGs act orthogonally to predict immunotherapy response or resistance, as we have previously shown^25^. Thus, our data suggest that ISGs have counterbalancing and temporally segregated functions with a key feature of ISG-mem genes being to directly contribute to ICB resistance.

Besides contributing to ICB-resistance, ISG-mem and related genes may also serve to ensure tonic activation of the RLR and IFN-I pathway. For example, ISG-mem genes include *Dhx58* and *Irf7* that are components of the RLR pathway^38^, as well as *Oas1* that can amplify autocrine IFN-I^25^. Growing evidence indicates that in multiple normal tissue compartments and cancer types, RLR and IFN-I signaling can control stem and progenitor cell differentiation dynamics, contributing to disease pathogenesis, adaptation to inflammation, or therapy resistance. In normal tissues, IFNs can have varying effects on hematopoietic stem cell (HSC) proliferation and survival, self-renewal, and differentiation bias particularly after infection and inflammation^53,54^. Similarly, RLRs can support developmental HSC formation, possibly through sterile inflammation by EREs^55^. Proliferation and differentiation of intestinal epithelial progenitors can also be controlled by IFNs and RLRs, an effect that can accelerate tissue regeneration after chronic inflammation-induced damage^56^. In CD8 T cells, IFN-I can drive differentiation of CD8 T cell progenitors toward exhaustion rather than effector-memory lineages, helping to prevent immune-mediated pathology during chronic infections^26^. In cancer, we show that ISG-mem^+^ RES_pre_ cells act as precursors that give rise to resistant, and possibly sensitive, epigenetic states that, like normal tissue stem/progenitor cells, is controlled by RLR and IFN-I signaling. Similarly, resistant cancer cells emerging from a single-cell-derived population after treatment with the TKI vemurafenib can exhibit precursor-like behavior and give rise to diverse features and phenotypes, including cells enriched in ISGs and RLRs^30^. In prostate cancer, subpopulations of cancer cells refractory to anti-androgen therapy can exhibit JAK/STAT-dependent lineage plasticity important for resistance^57^. Collectively, these findings suggest that in normal tissues and cancer cells, minor subpopulations with progenitor-like properties can utilize RLR and/or IFN-I signaling to alter cell fate decisions brought on by chronic inflammation or therapy pressure.

Lineage tracing analysis of our ICB-relapsed tumor model reveals extensive transitions between ISG-mem^+^ RES_pre_ precursor cells and cancer cells with either sensitive or resistant chromatin states. However, after ICB activates an anti-tumor immune response, survival of lineages primarily comprised of resistant Epi-Res2 chromatin states are favored. We show that pharmacologically targeting RLR and IFN-I signaling using JAKi + TBK1i prevents differentiation into this resistant chromatin state and skews the composition heavily toward sensitive states, resulting in ICB responsiveness irrespective of lineage composition prior to drug therapy. Details as to why Epi-Res2 chromatin states are resistant to ICB will require additional investigation. As mentioned, ISG-mem genes and additional genes controlled by cis-REs of ISG-mem and Epi-Res2 are likely key; however, other mechanisms may contribute or might be under the influence of these cis-REs. For example, cell populations like ISG-mem^+^ RES_pre_ with precursor-like properties are necessarily less differentiated and may share properties with tumor-initiating cells. Cytokine-driven dedifferentiation can promote immunotherapy resistance due to loss of differentiation antigens^58,59^, and cancer cells with tumor-initiating properties can exhibit low-levels of antigen presentation^60^, express TGF-beta-responsive inhibitory molecules^61^, or downregulate activating receptors for NK cells^62^. Thus, some of these other immune evasion mechanisms attributed to de-differentiated cancer cells may play a role in ICB resistance brought on by chronic IFN and inflammatory memory. Alternatively, epigenetic states such as Epi-Res1 that do not appear related to inflammatory memory may represent additional unidentified mechanisms of acquired resistance.

We have recently reported promising clinical and immune responses by using JAK inhibitors with anti-PD1 for first-line metastatic NSCLC^15^, results that have been independently observed in the setting of anti-PD1-refractory Hodgkin lymphoma^16^. In our NSCLC study, we show that using JAKi to interfere with inflammatory signaling, including IFN-I, is associated with tumor response and appears to inhibit the ability of IFN-I from directing precursor/progenitor CD8 T cells toward terminal and exhaustion lineages. However, although response rates were high, a subset of patients nonetheless did not experience partial or complete response. These non-responders were characterized by high baseline inflammation that was refractory to JAKi. We now show that JAKi alone is insufficient to effectively abrogate features of inflammatory memory acquired by ICB-resistant cancer cells. In particular, effective reversal of the enhanced chromatin accessibility of resistance-associated activated enhancers and of EREs that are part of a MDA5 positive feedback loop require both JAKi and TBK1i, consistent with the need to co-delete STAT1 and IRF3 to abrogate epigenetic changes of inflammatory memory. Accordingly, JAKi + TBK1i was markedly more effective than JAKi alone or TBK1i alone at preventing ICB relapse in mouse tumor models. It has been reported that TBK1i alone can improve ICB^63^; however, we surmise that in the context of inflammatory memory and acquired resistance, poised enhancers and positive feedback circuits set in place by inflammatory memory lowers the threshold for gene activation, making inhibition by JAKi or TBK1i alone difficult to achieve. Thus, particularly for patients with high levels of inflammation related to inflammatory memory, the combination of JAKi + TBK1i may hold promise as an approach to inhibit acquired ICB resistance by building on recent strategies that target persistent inflammation and IFN signaling. Given the manifold diseases and conditions that may involve inflammatory memory – autoimmune diseases, aging, neurodegeneration, COVID – therapeutic approaches relevant for inflammatory memory may extend beyond cancer.

## ACKNOWLEDGMENTS

A.J.M., R.A.G., J.S., and N.R.Z. are supported by The Mark Foundation for Cancer Research. A.J.M. and D.Y. are also supported by the Parker Institute for Cancer Immunotherapy. A.J.M. is additionally supported by the Breast Cancer Research Foundation. We thank David Feldser for sharing the KP cell line utilized throughout this study, and Andrew Modzelewski for insightful discussion on endogenous retroelement biology.

## AUTHOR CONTRIBUTIONS

Conceptualization, J.Q., D.Y., and A.J.M.; Primary Investigation, D.Y. and N.D.; Investigation Support; B.Y., Y.S., S.W., D.R., and Y.X.; Methodology, D.Y. and J.Q.; Resources, J.C.; Formal Analysis, J.Q., X.E.C., and A.J.M.; Software, J.Q., X.E.C., T.Z., and V.N.; Visualization, J.Q., X.E.C., and A.J.M.; Writing, J.Q. and A.J.M.; Supervision, A.J.M., R.A.G., J.S., and N.R.Z.; Project Administration, A.J.M.

## SUPPLEMENTAL FIGURE LEGENDS

**Figure S1.**
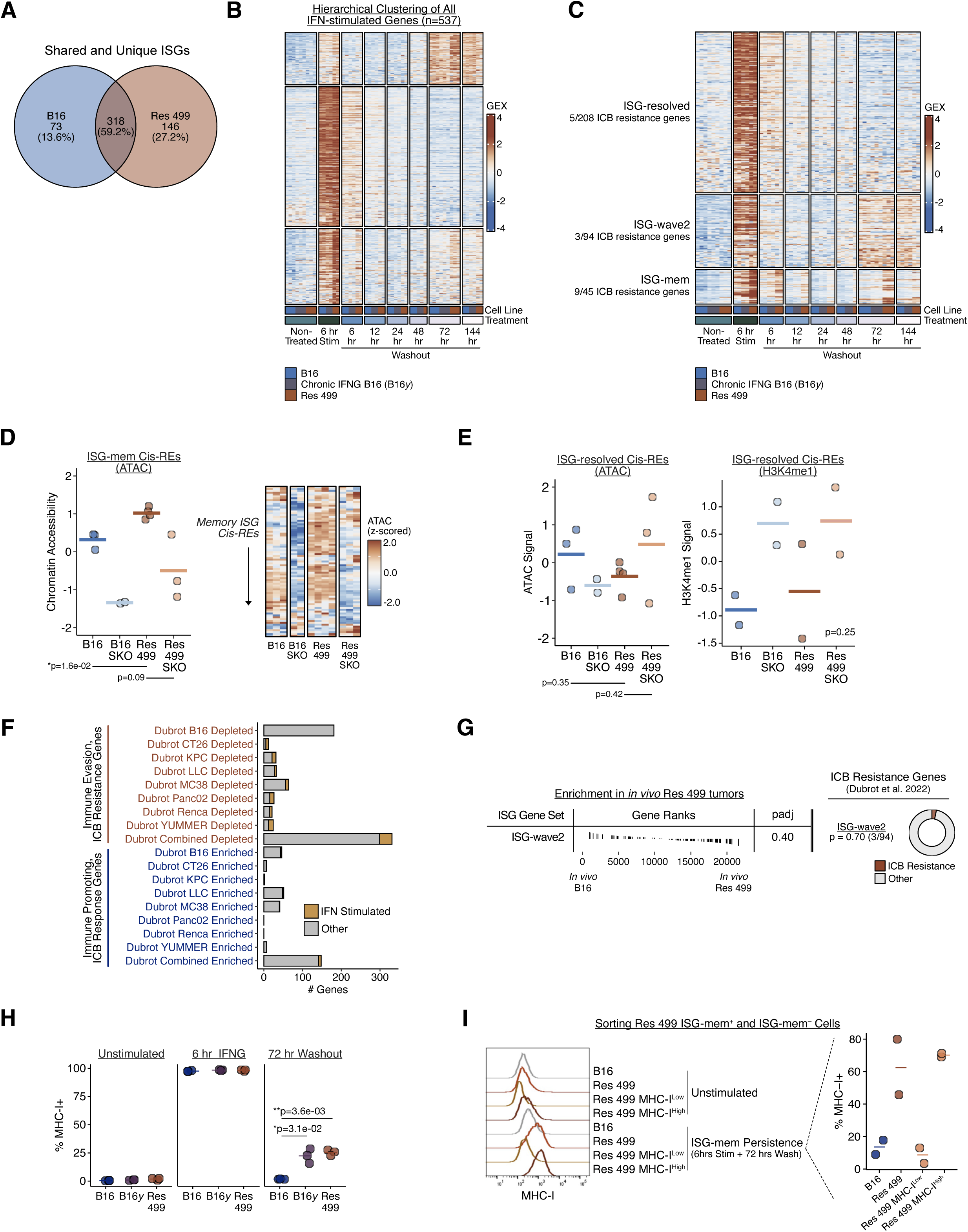
Features of memory ISGs associated with immunotherapy resistance (Related to Figure 1). **A)** Venn diagram displaying the number of shared and unique ISGs in B16 and Res 499 cells, identified from the IFN stimulation and washout experiment described in Figure 1B. **B)** Scaled gene expression of all B16 and Res 499 ISGs from (A) are shown for the indicated cell lines. Hierarchical clustering was performed to identify temporal expression patterns. **C)** Scaled gene expression of manually curated Res 499 ISGs that fall into three main temporal expression patterns (Resolved, Wave 2, Memory) are shown for the indicated cancer cell lines. **D)** Summary GSVA enrichment (left) and heatmap (right) of chromatin accessibility scores for memory ISG cis-REs in B16 or Res 499 cells with or without STAT1 KO (SKO) sorted from *in vivo* tumors (left; n=3 per condition). **E)** Summary chromatin accessibility (left) and H3K4me1 (right) GSVA enrichment scores of resolved ISG cis-REs in B16 or Res 499 cells with or without STAT1 KO (SKO) sorted from *in vivo* tumors (n=2-3 per condition). P-value for H3K4me1 samples was determined by a one-side ANOVA test. **F)** The number of genes depleted (i.e., ICB resistance genes) or enriched identified by Dubrot et al. genome-wide *in vivo* CRISPR screening from each indicated syngeneic mouse tumor model. The proportion of genes that are B16 and/or Res 499 ISGs from (A) are indicated in orange. **G)** Gene set enrichment analysis (GSEA) for ISG-wave2 on genes ranked by differential expression between Res 499 and B16 *in* vivo tumors (left). Over-representation analysis results for enrichment of Dubrot et al. ICB-resistance genes in ISG-wave2 (right). P-values determined by Fisher’s exact test. **H)** Normalized percent MHC-I^+^ cells for the indicated cell line at baseline, after 6 hours of IFNG stimulation, or at 72 hours washout. **I)** Representative flow cytometry histogram of MHC-I at baseline (unstimulated) or at 72 hours washout after IFNG stimulation for B16, Res 499, or Res 499 MHC-I^Low^ or Res 499 MHC-I^High^ previously sorted from a persistent ISG-mem *in vitro* assay (Fig. 1J). Summary plot of percent MHC-I^+^ for the indicated cell line at 72 hours washout after 6 hours IFNG stimulation is also shown (right). Unless otherwise indicated, P-values for two sample comparisons were determined by a two-sided t-test.

**Figure S2.**
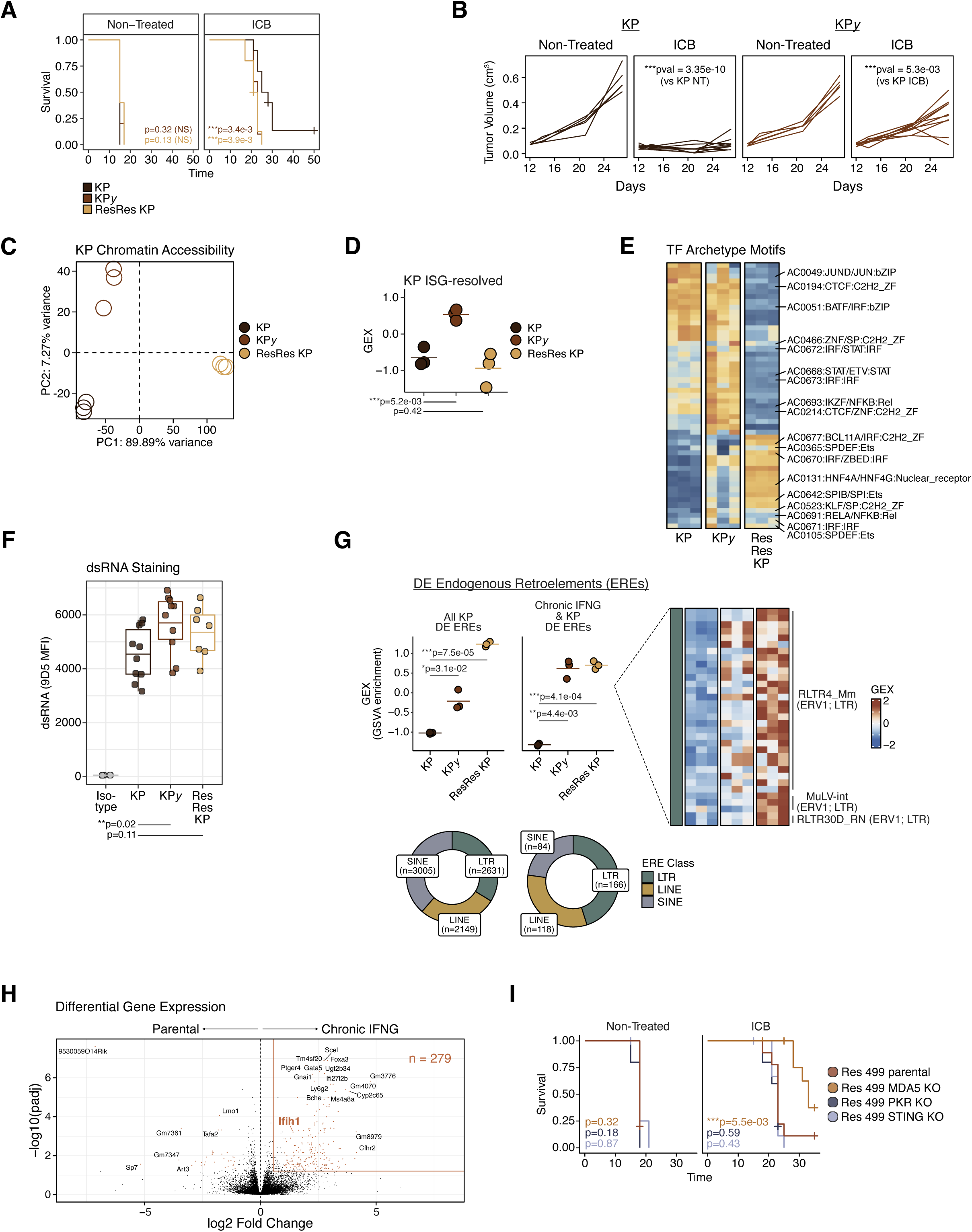
Inflammatory memory, chronic virus mimicry, and ICB-resistance features for lung cancer model (Related to Figure 2). **A)** Survival of mice with KP lung cancer, KP*y*, or ResRes KP tumors either non-treated (n=5 per condition) or treated with anti-CTLA4 + anti-PD1/PDL1 + CD40 agonist (ICB) (n=10 per condition). **B)** Tumor growth from mice with KP or KP*y* either non-treated (n=5 per condition) or treated with anti-CTLA4 + anti-PD1/PDL1 + CD40 agonist (ICB) (n=10 per condition). **C)** Principal component analysis of genome-wide chromatin accessibility profiles from KP parental, KP*y*, and ResRes KP cancer cells sorted from *in vivo* tumors. **D)** Summary GSVA enrichment score for expression of ISG-resolved in cancer cells sorted from the indicated *in vivo* tumor (n=3 per condition). **E)** ChromVAR chromatin accessibility scores of the top 20 variable TF archetype motifs and STAT/IRF motifs of interest in the indicated *in vivo* KP tumor. Columns are individual biological replicates. **F)** MFI of dsRNA levels using the 9D5 antibody for the indicated cancer cells sorted from *in vivo* tumors (n=6-9 per condition). **G)** Summary GSVA enrichment score for expression of EREs with significantly increased differential expression (DE) in ResRes KP or both KP*y* and ResRes KP cells sorted from *in vivo* tumors (n=3 per condition). Individual biological replicates grouped in the same order as the summary plot are shown in the heatmap (right). ERE class composition is indicated in a donut plot (bottom). **H)** Volcano plot of log2 fold changes in gene expression and -log10 P-values for differential gene expression analysis comparing cancer cells sorted from parental KP and LLC1 tumors versus tumors from chronic IFNG treated cells. **I)** Survival of mice with Res 499 tumors or tumors from Res 499 cells with the indicated PRR KO either non-treated (n=5 per condition) or treated with anti-PDL1 + anti-CTLA4 (ICB) (n=10 per condition). P-values for two sample comparisons were determined by a two-sided t-test. P-values for survival were determined by a log-rank test, and for tumor growth by a mixed-effect regression model.

**Figure S3.**
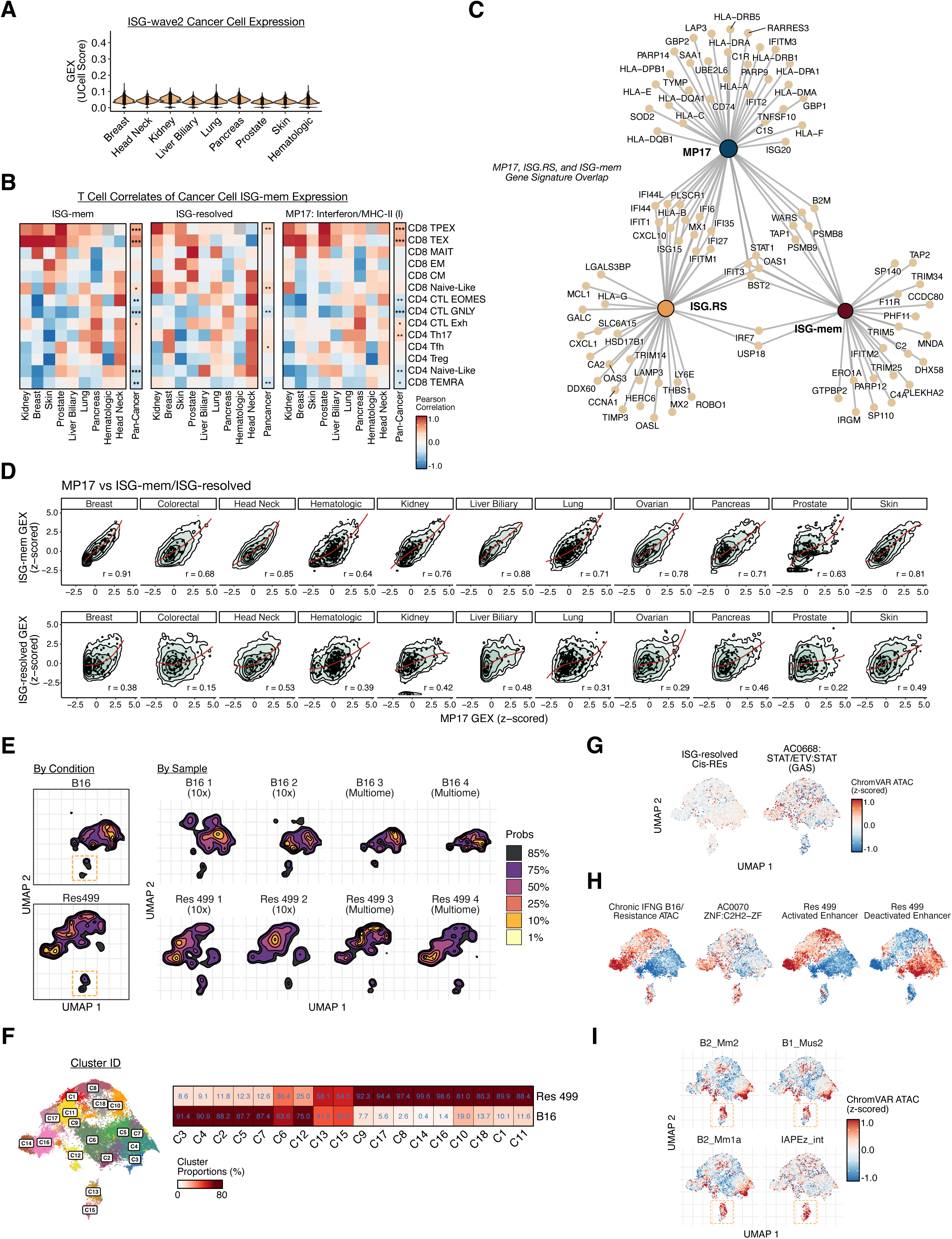
Features of cancer cell subpopulations expressing memory ISGs from human and mouse tumors (Related to Figure 3). **A)** Expression of ISG-wave2 in cancer cells for the indicated cancer type. Gene expression is the single-cell UCell score. **B)** T cell correlates for the indicated ISG signatures. Color values represent Pearson correlations between mean cancer cell ISG expression and proportion of T cell subsets (rows) for the indicated cancer type (columns). P-values were determined by a pan-cancer correlation significance test with significant correlations denoted with an asterisk. **C)** Overlap of individual genes from human MP17, ISG.RS, and mouse memory ISG signatures. **D)** Correlation between MP17 expression and expression of ISG-mem (top row) or ISG-resolved (bottom row) in cancer cells from the indicated cancer type. The highest density regions at 99%, 95%, 80%, 50%, and 25% are plotted. A spline regression line is fit through the data and the Spearman correlation is shown. **E)** Density plot of UMAP embedding of chromatin accessibility features for biological replicates of B16 and Res 499 cells sorted from *in vivo* tumors. ISG-mem^+^ RES_pre_ population is highlighted with an orange box. **F)** UMAP embedding of chromatin accessibility features for B16 and Res 499 cells colored by SnapATAC cluster ID (left). Proportion of cells in each SnapATAC cluster for each indicated condition is shown in the heatmap (right). **G-I)** ChromVAR chromatin accessibility scores (bias-corrected deviation scores) for the indicated archetype TF motif, chromatin signature, or chronic IFNG-induced ERE subfamily overlaid on the UMAP embedding from Fig. 3F. *p < 0.05; **p<0.01; ***p<0.001.

**Figure S4.**
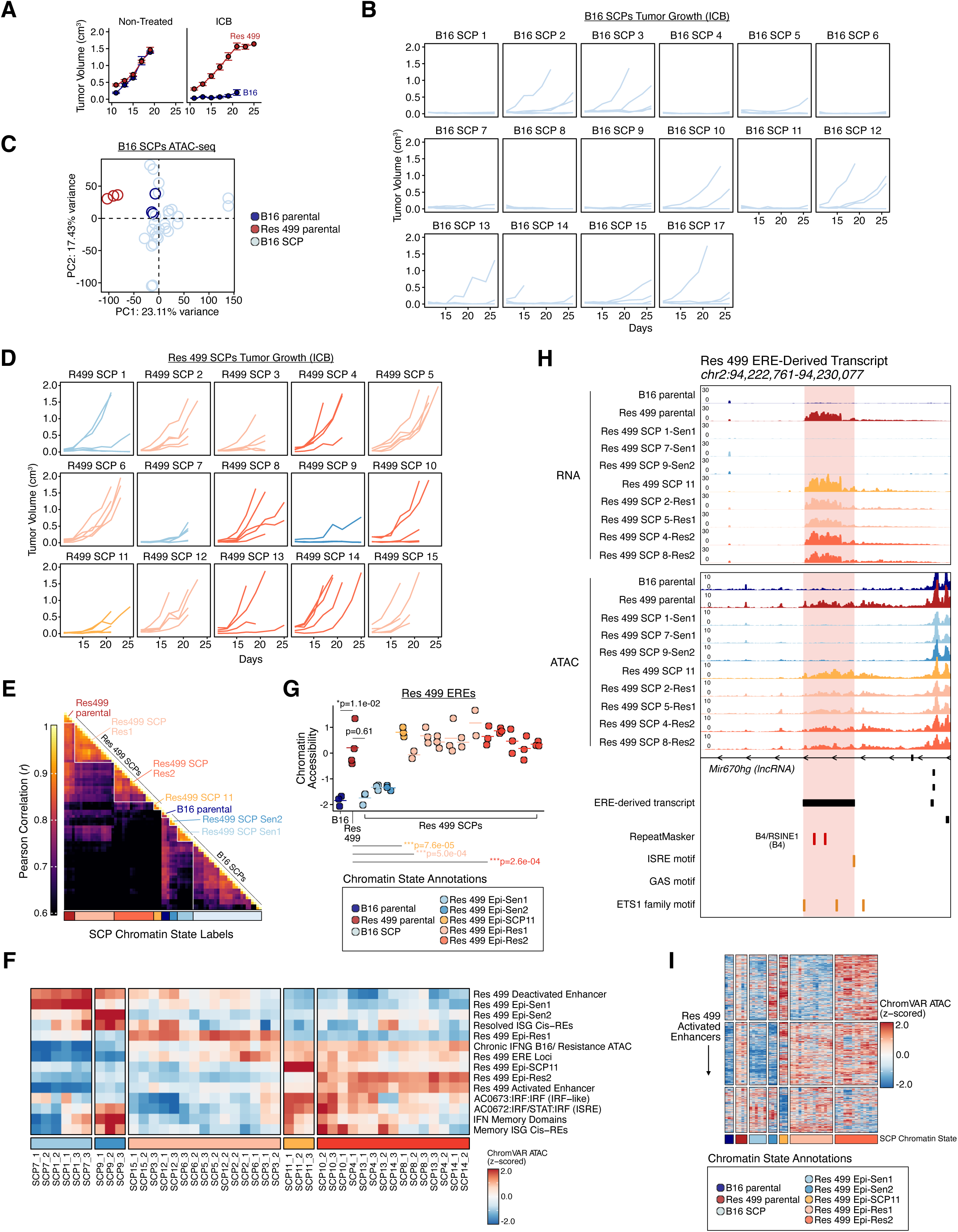
Features of B16 and Res 499 SCPs (Related to Figure 4). **A)** Tumor growth of mice with B16 and Res 499 parental tumors and **B)** B16 SCPs treated with anti-CTLA4 + anti-PDL1 (ICB) (Non-Treated, n=5 per condition; ICB, n=5-10 per condition). **C)** Principal component analysis of genome-wide chromatin accessibility profiles from B16 SCPs cells sorted from *in vivo* tumors. **D)** Tumor growth of mice with Res 499 SCPs treated with anti-CTLA4 + anti-PDL1 (n=5-10 per condition). SCPs are color-coded by SCP chromatin state as shown in the legend for (G). **E)** Hierarchically clustered pairwise Spearman correlations of chromatin accessibility profiles between all B16 and Res 499 SCP cells sorted from *in vivo* tumors. Bottom annotation bar depicts the SCP chromatin state used to group SCPs, as indicated by the legend for (G). **F)** ChromVAR chromatin accessibility scores (bias-corrected deviation scores) of the indicated chromatin signatures (rows) from Res 499 SCPs sorted from *in vivo* tumors and grouped by SCP chromatin state shown in the legend for (G). **G)** Summary ATAC GSVA enrichment for chromatin accessibility scores of Res 499-activated EREs for B16, Res 499, and SCPs sorted from *in vivo* tumors (n=3 per condition). SCPs are color-coded by chromatin state shown in the legend. **H)** Representative RNA and chromatin accessibility (ATAC) tracks at an ERE-derived transcript loci from the indicated cancer cells sorted from *in vivo* tumors. ERE repeat pairs are annotated in red, TF motifs are annotated in orange. **I)** Chromatin accessibility of Res 499 activated enhancers for B16, Res 499, and SCPs sorted from *in viv*o tumors (n=3 per condition). SCPs are grouped by chromatin state shown in the legend. P-values for two sample comparisons determined by a two-sided t-test. Error bars represent standard errors.

**Figure S5.**
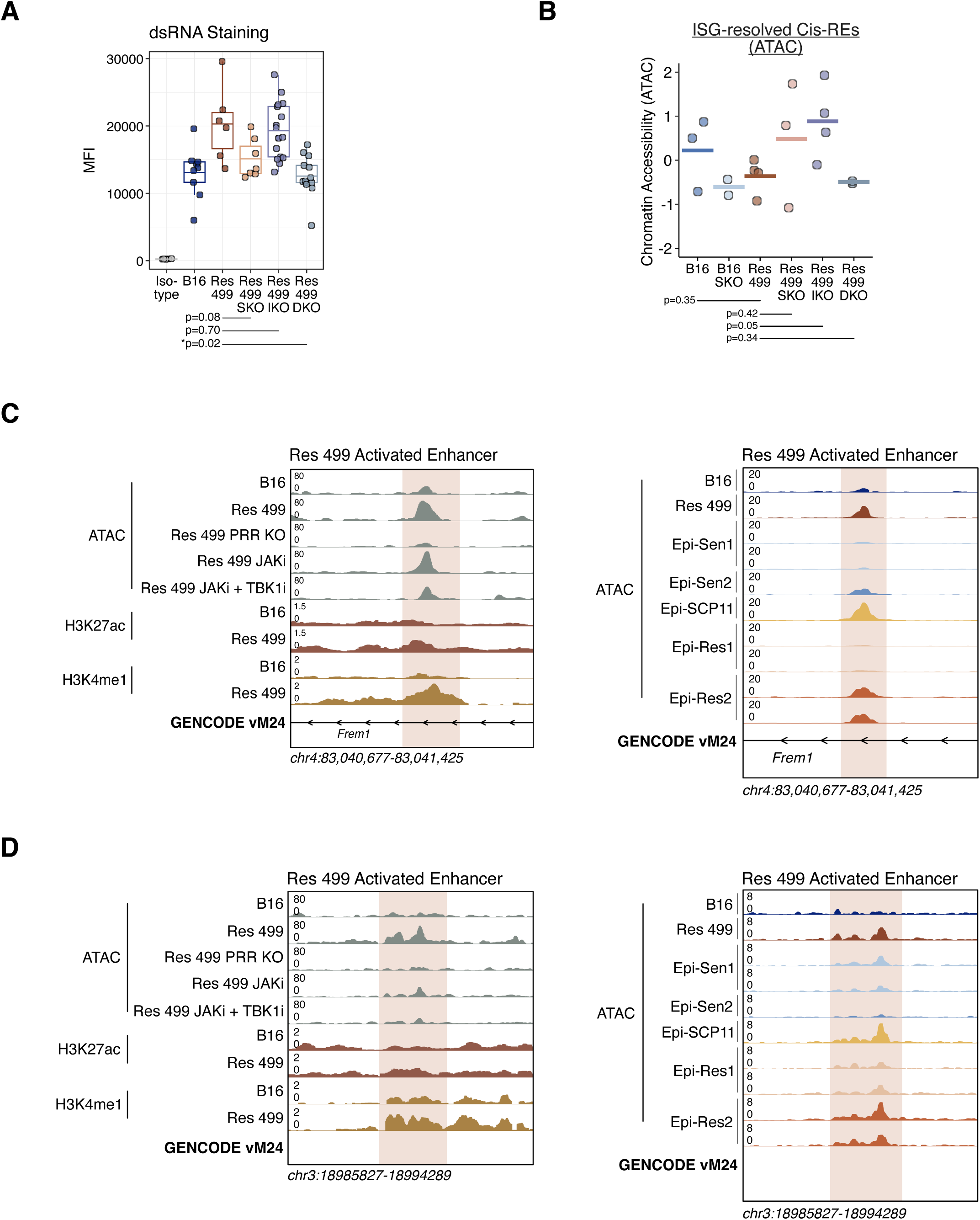
Effect of STAT1, IRF3, and JAKi plus TBKi combination therapy on dsRNA levels and epigenetic features of inflammatory memory (Related to Figure 5). **A)** Mean fluorescence intensity (MFI) of dsRNA levels using the 9D5 antibody in B16 or Res 499 cells with or without STAT1 KO (SKO), IRF3 KO (IKO), or STAT1 and IRF3 KO (DKO) sorted from *in vivo* tumors (n=6-16 per condition). **B)** Summary chromatin accessibility GSVA enrichment score of ISG-resolved Cis-REs for B16 or Res 499 cells with or without STAT1 KO (SKO), IRF3 KO (IKO), or STAT1 and IRF3 KO (DKO) sorted from *in vivo* tumors (n=2-4 per condition). **C)** Representative chromatin accessibility (ATAC) and CUT&RUN tracks for H3K27ac and H3K4me1 for a Res 499 activated enhancer loci (same as in Fig. 5G) for B16 or Res 499 cells expanded *in vitro* with or without JAKi +/- TBK1i and sorted from *in vivo* tumors (left). For comparison, Res 499 cells with MDA5/RIG-I KO (PRR KO) were used. Also shown are representative chromatin accessibility tracks at the same enhancer loci for Res 499 SCP 11 (color-coded orange) and other Res 499 SCPs with the indicated SCP chromatin state and sorted from *in vivo* tumors (right). **D)** Same as (C) but for another Res 499 activated enhancer loci. GENCODE vM24 gene annotations are shown. P-values for two sample comparisons were determined by a two-sided t-test.

**Figure S6.**
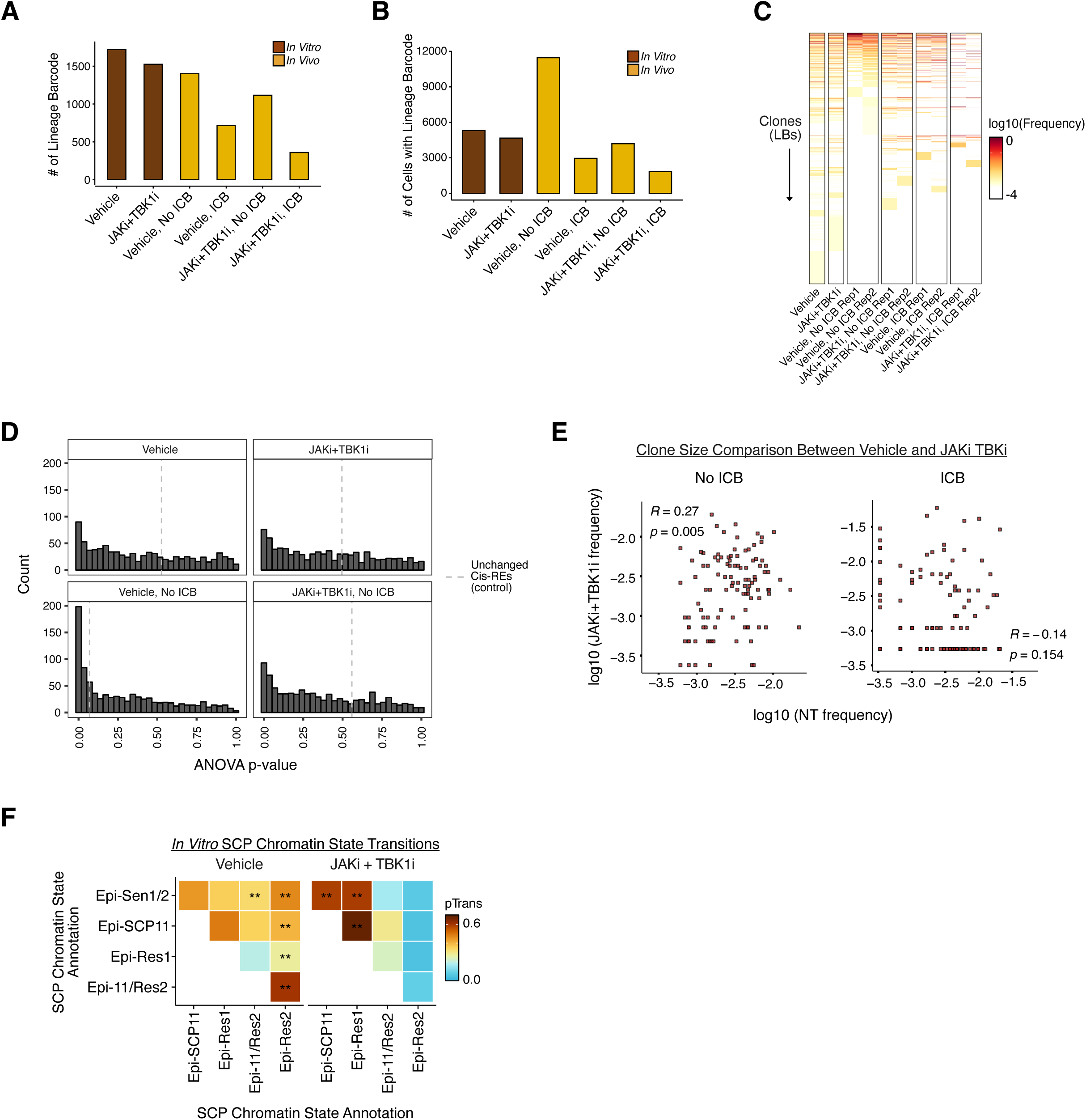
Lineage barcoding of Res 499 cancer cells to track clonal fate and composition (Related to Figure 6). **A)** Number of unique lineage barcodes or **B)** number of cells with detected lineage barcode after *in vitro* culture of Res 499 cells with or without JAKi + TBK1i (brown) or after *in vitro* culture with or without JAKi + TBK1i followed by implantation into mice treated with or without ICB (dark yellow). See Fig. 6A for experimental design. **C)** Heatmap of clonal frequency for cells with each lineage barcode (rows) for the same indicated conditions. **D)** Histograms of P-values from ANOVA tests for identifying heritable chromatin features. Dashed lines indicate the P-values of unchanged cis-REs. **E)** Clonal frequency of each lineage barcode comparing *in vitro* vehicle control (x-axis) versus *in vitro* JAKi + TBK1i treatment (y-axis) examined after implantation into mice (left), or after implantation into mice followed by ICB treatment (right). **F)** Pairwise transition-index values (pTrans-index) between Res 499 cells annotated by the indicated SCP chromatin state after *in vitro* culture with vehicle or with JAKi + TBK1i. Higher pTrans-index values indicate greater barcode sharing or developmental relatedness. Values that are significantly higher in one treatment condition compared to the other are marked with an asterisk. *p < 0.05; **p<0.01; ***p<0.001.

**Figure S7.**
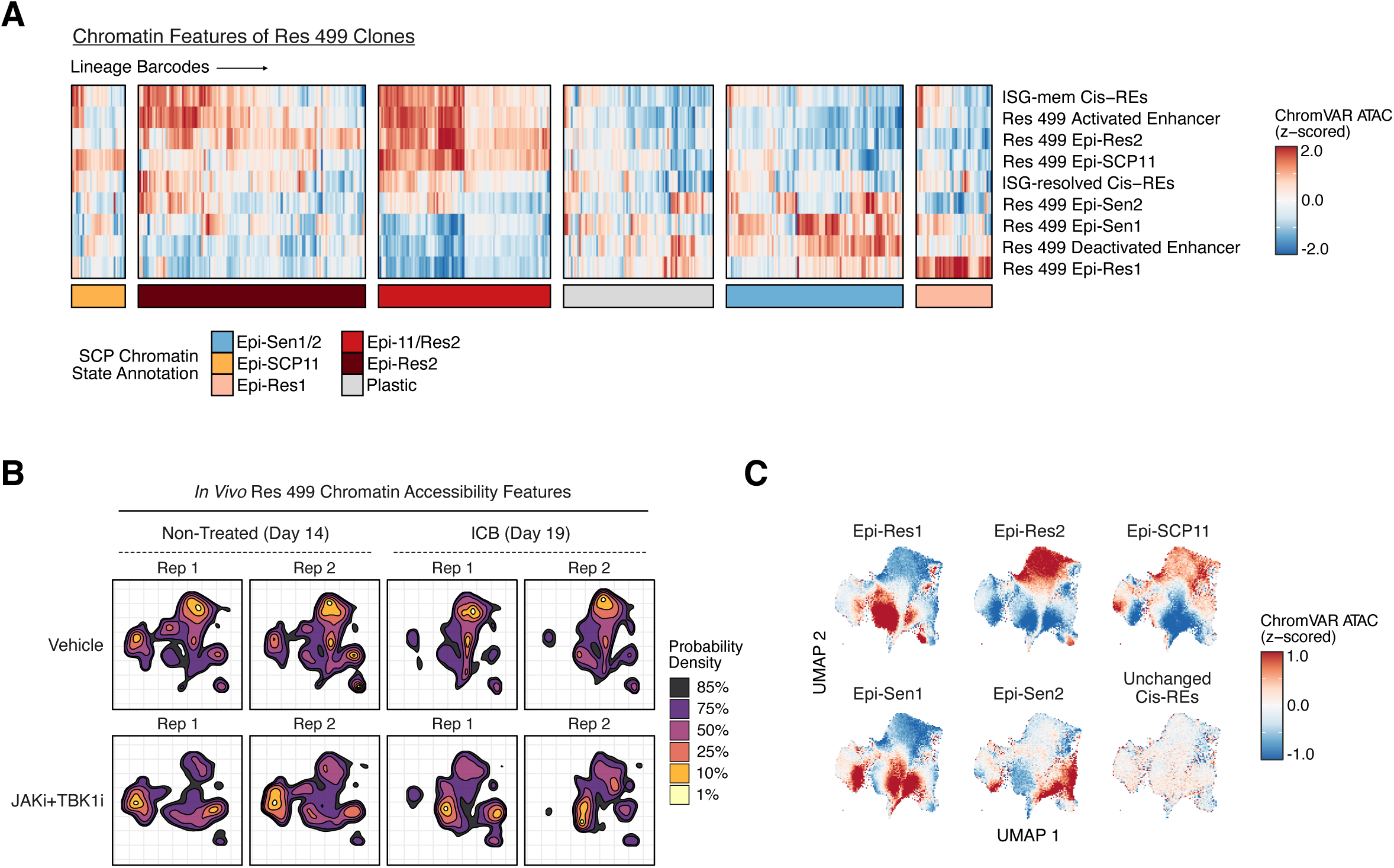
Effect of JAKi plus TBK1i on survival of Res 499 clones after tumor implantation and treatment with ICB (Related to Figure 7). **A)** Mean scaled ChromVAR chromatin accessibility scores for the indicated chromatin signature (rows) for clones annotated by its dominant SCP chromatin signature. Plastic annotation indicates clones without a dominant SCP chromatin signature. **B)** Density contour plot of UMAP embedding for chromatin accessibility features for Res 499 cells expanded *in vitro* with vehicle or with JAKi + TBK1i, followed by implantation into mice that were either non-treated or treated with ICB. **C)** ChromVAR chromatin accessibility scores (bias-corrected deviation scores) of Res 499 SCP chromatin signatures plotted overlaid on a UMAP embedding from (B).

## EXPERIMENTAL PROCEDURES

### Mice

All animal experiments were performed according to protocols approved by the Institutional Animal Care and Use Committee of the University of Pennsylvania. Five-to seven-week-old female C57BL/6 (stock# 027) and BALB/c (stock# 28) were obtained from Charles River Laboratory. Mice were maintained under specific pathogen free conditions and randomly assigned to each experimental group.

### In vivo mouse studies

Tumor cells were subcutaneously injected into flanks of mice. The number of cells injected varied from 50,000 to 100,000 cells depending on the cell line. Antibody treatments were carried out on Days 5, 8 and 11. Antibodies were from Bio X Cell: anti-PD1 (RMP1-14), anti-PD-L1 (10F.9G2), anti-CTLA4 (9H10).

### Flow cytometry and cell sorting

Two weeks following tumor injection, tumors were harvested, digested with collagenase IV (EMSCO/Fisher) for 30 minutes at 37 degrees C. The suspension was filtered, treated with ACK lysis buffer (Quality Bio) for 5 minutes on ice. Staining was carried out first in Live/dead fixable aqua dead cell staining kit (Thermo Fisher Scientific) and TruStain fcX (Biolegend) for 15 minutes at 4 °C, followed by surface antibody staining for 30 minutes at 4 °C. Intracellular staining was carried out using Foxp3/Fix/Perm kit (Thermo Fisher Scientific). Cell sorting was carried out on an Aria cell sorter when necessary. Fluorescently labeled anti-mouse antibodies against CD45 (104) and H-2Kb/H-2Db (28-8-6) were obtained from BioLegend.

### ATAC and RNA sequencing

Single cell suspension was obtained as described above from Day 15 tumors and tumor cells were sorted by gating on live/dead aqua negative and CD45 negative cells. 50,000 or 100,000 cells were sorted for ATAC-seq or RNA-seq respectively. RNA was extracted using Direct-zol RNA kits (zymo research) with on column DNase I treatment. Library was prepared using Truseq Stranded Total RNA Library Prep Gold kit. ATAC-seq libraries were prepared as described previously^39^. Both libraries were sequenced on NovaSeq 6000 with 50 bp paired-end reads.

### Cytoplasmic dsRNA Staining

*In vivo* harvested mouse tumors were stained with L/D Aqua (1:600) and CD45-AF700 (1:200) in a 96-W plate on-ice for 15-20 minutes, then fixed with 2% PFA/PBS for 15 minutes on ice. Cells were permeabilized with 0.1% Saponin/FACS buffer for 15 minutes, stained with 9D5 dsRNA rabbit IgG mAb (1:500) in 0.1% Saponin/FACS Buffer for 15 minutes on ice, washed with saponin buffer, stained with Donkey aRabbit IgG-PE (1:400) in 0.1% Saponin/FACS Buffer for 15 minutes on ice, washed with saponin buffer, resuspended in FACS buffer, then flowed on an LSR to assay for dsRNA on live/CD45-cancer cells.

### *In vitro* Persistent Memory ISG Assay

Cells were plated at 22,000 cells/W in a 24-W plate overnight. Cells were then stimulated with mIFNg (100ng/mL) for 6 hours, then washed twice with complete media and then replenished with complete media. 72 hours later, cells were collected via trypLE treatment, stained with MHC1-AF647 (1:200) in FACS buffer (2% FBS/ PBS) on ice for 15-20 minutes, then readout on an LSR for persistent MHC-I expression.

### Generation of Watermelon library barcoded Res499 cells

10,000 Res499 cells were spin transduced with Watermelon library barcode virus at an MOI of ∼0.22 at 2225 RPM for 30 minutes at 32 degrees C with 8ug/mL polybrene in a 48-W plate. 24 hours later, lentivirus transduced media was replaced with complete media, and cells were expanded for 3 days, then transferred to a 6-W plate and left expanded for 5-6 additional days. Following expansion, cells were then collected as a single-cell suspension and sorted for GFP+ cells on an ARIA sorting machine.

### *In vitro* Treatment of Res499 cells with 3 weeks JAKi + TBK1i

Res499 cells were treated with Ruxolitinib (5uM) + Cmpd1 (1uM), or DMSO as vehicle (no more than 1:2000 dilution), and cells were split every other day, replenishing drug media, for 3 weeks, followed by 1 week washout with no drug. Following 1 week drug washout, cells were either frozen down or implanted directly into mice.

### RNA-seq processing

Adapters were trimmed with cutadapt version 2.9. Gene-based quantification was performed with salmon version 1.8.0 (--validateMappings --rangeFactorizationBins 4 --seqBias --gcBias). Salmon counts were tabulated using the R package *tximport* version 1.22.0. For the final count matrix, raw counts were normalized using the *vst* transformation from DESeq2 version 1.34.0.

### ATAC-seq processing

Sequencing reads were processed using the ENCODE ATAC-seq pipeline, part of the ENCODE Uniform Processing Pipelines series (https://github.com/ENCODE-DCC/atac-seq-pipeline). In brief, adapters were trimmed using cutadapt, and reads were aligned to the mm10 genome using Bowtie2. After alignment, duplicate reads and reads mapping to chrM were removed. To correct for the Tn5 offset, bam file reads were shifted (“+” stranded +4 bp, “-“ stranded -5 bp). Peak calling was performed with MACS2, and high-confidence peaks were filtered for using the irreproducible discovery rate (IDR), which selects for peaks that have consistent ranks across replicates. Overlapping peaks were merged across all conditions to generate a consensus peak set. A raw count matrix was generated by counting the number of insertions that overlapped each region in the consensus peak set. For the final count matrix, raw counts were normalized using the *vst* transformation from DESeq2 version 1.34.0.

#### Identifying Persistent IFN ISG Signatures

IFNG response genes were empirically identified using DESeq2 version 1.34.0 and hierarchically clustered by RNA expression over the persistent IFN time course. Three major temporal transcriptional patterns were revealed and further curated to robust gene signatures: “Resolved ISGs”, like canonical ISGs, are induced by IFNG and return to their unstimulated expression levels by 72 hours washout. “Wave 2” ISGs exhibit diminished expression in the 6-24 hour washout period, but undergo a second wave of transcription at 72 hours. “Memory ISGs” exhibit a significantly augmented acute induction and persistent expression in Res 499 and B16y at 6 hours stimulation and 72 hours washout, respectively.

### Identifying peak-to-gene links

To identify putative cis-regulatory elements for individual genes, we fit multiple linear regression models using chromatin accessibility features as predictors and the RNA expression of each gene of interest as the dependent variable. For each gene of interest, we included as features all ATAC-seq peaks within a 184 Kb cis-regulatory window, the median size of a mouse topological associated domain. Peaks with non-zero coefficient estimates and associated with genes with a variance explained of R^2^ > 0.4 were considered putative cis-regulatory elements (cis-REs).

### Identifying differentially expressed endogenous retroelements (EREs)

RNA-seq reads were aligned using STAR version 2.7.1a with custom parameters allowing for multi-mapped reads (--outFilterMultimapNmax 100 --winAnchorMultimapNmax 200 -- outFilterMismatchNoverLmax 0.04 --alignSJoverhangMin 8 --alignSJDBoverhangMin 1 -- alignIntronMin 20 --alignIntronMax 1000000 --alignMatesGapMax 1000000). ERE subfamilies were quantified using TEtranscripts version 2.2.1 in *multi* mode^64^, which uses an expectation maximization algorithm to assign multi-mapped reads. ERE loci were quantified using squire Count version *0.9.9.92*^65^. We generated normalized gene + ERE count matrices and identified differentially expressed ERE subfamilies or loci using DESeq2 version 1.34.0.

### scATAC-seq processing

10X scATAC-seq data was demultiplexed and aligned to the mm10 genome using CellRanger ATAC 2.0.0 and default parameters. We processed the data for downstream analysis using SnapATAC^66^. Paired-end reads from the 10X-generated bam file were converted into fragments and only fragments with MAPQ > 30, properly paired, and 50-1000 bp in length were retained. Cell barcodes with >1000 unique fragments and a promoter ratio (percentage of fragments overlapping annotated promoter regions) between 0.2 and 0.8 were retained. We binned the genome into 5000 bp windows (rather than peaks) and counted insertions within each bin. Bins overlapping ENCODE blacklist regions, mitochondrial DNA, or in the top 5% of invariant features were filtered out. We binarized the bin-by-cell matrix and generated a normalized Jaccard similarity matrix. We used SnapATAC’s implementation of dimensionality reduction with eigenvector decomposition, chose 14 eigenvectors for downstream analysis, and visualized cells using uniform manifold approximation and projection (UMAP). Harmony was used to remove batch effects across different scATAC-seq runs. We called peaks for individual communities of cells (Louvain clusters) in a k-nearest neighbor graph and called accessible chromatin peaks for each cluster. Overlapping peaks were merged across all conditions to generate a consensus peak set. We then counted insertions within each peak to generate a peak-by-cell matrix, which we used as input for chromVAR to calculate TF motif deviations and enrichment of signatures of interest.

### scATAC-seq pseudotime analysis

Since Res 499 is a direct relapse of B16, we sought to infer putative lineage trajectories using Cicero^41^. In brief, we generated a CDS object using a count matrix and consensus peak set generated by SnapATAC, performed dimensionality reduction using latent semantic indexing (LSI), learned a trajectory graph, and ordered cells rooting pseudotime in the cluster of cells with the lowest chromatin accessibility at Res 499 activated enhancers.

### Motif Analysis

To avoid conflating TFs with highly similar motifs, we used the 693 non-redundant position weight matrices for “archetype motifs”^67^ from clustering over 5000 motif PWMs from JASPAR, CIS-BP, TRANSFAC, HOCOMOCO, and UniProbe (https://resources.altius.org/#jvierstra/projects/motif-clustering-v2.0beta/). ISRE motifs were represented by AC0672:IRF/STAT:IRF, GAS motifs were represented by AC0668:STAT/ETV:STAT, NFKB RELA motifs were represented by AC0693:IKZF/NFKB:Rel, and ETS1 binding sites were represented by AC0641:ETV/ETS:Ets. We used these PWMs to calculate GC-bias corrected motif accessibility deviations in bulk and single-cell ATAC-seq data using ChromVAR.

### ERE subfamily enrichment in scATAC-seq data

Genomic regions for each ERE subfamily were extracted from *mm10* RepeatMasker. ChromVAR version 1.16.0^68^ was used to calculate chromatin accessibility enrichment at each ERE subfamily signature in single-cell ATAC-seq data.

### Human tumor scRNA-seq analysis

scRNA-seq datasets were downloaded from the Curated Cancer Cell Atlas (available datasets as of August 2023)^39^ (https://www.weizmann.ac.il/sites/3CA/) **(Supplementary Data Table 1)**. Malignant and T cells were subsetted using previously determined annotations curated by Gavish et al. and only samples comprising over 50 malignant cells and over 50 T cells were retained. The final pan-cancer dataset analyzed was composed of 32 individual studies and 201 tumor samples. Gene expression counts of malignant cells were log normalized and UMAPs were generated separately for each sample using Seurat version 4.3.0. Ucell single-cell expression scores, which are robust to dataset size and heterogeneity, were calculated using Ucell version 1.3.1^69^ for cancer cell meta-programs and ISG gene signatures. We determined T cell subtype annotations by mapping T cells onto a human tumor-infiltrating CD8+ lymphocyte atlas [https://github.com/ncborcherding/utility] using ProjectTILs version 3.1.2^70^. Mean ISG expression in malignant cells and proportion of each T cell subtype was calculated for each tumor sample.

### Assigning SCP chromatin states to single cells

Res 499 SCP ATAC signatures were derived by one vs. all differential accessibility analysis using edgeR (p < 0.05, log2fc > 1). Highly similar single cells were aggregated for each sample separately using SEACells version 0.3.3 for scATAC-seq data^71^ (approximately 75 cells per SEACell). Each SEACell was considered to be enriched for an SCP chromatin state or Res 499 enhancer signature if its cells’ z-scored ChromVAR deviation scores were significantly larger than 0 (one-sample t-test). Individual cells were then assigned to an SCP chromatin state using the following criteria:

1. SEACell enriched for sensitive SCP signatures and Res 499 deactivated enhancers were annotated as Epi-Sen1/2.
2. SEACells enriched for resistant SCP signatures and Res 499 activated enhancers were annotated as Epi-Res1 or Epi-Res2.
3. SEACells enriched for the SCP 11 signature and no other SCP signatures were annotated as Epi-SCP11.
4. SEACells enriched for both Epi-SCP11 and Epi-Res2 were annotated as hybrid Epi-11/Res2.
5. SEACells enriched for conflicting SCP signatures were annotated as Plastic.

### Identify heritable chromatin features

To identify heritable chromatin features, we performed Analysis of variance (ANOVA) test across clones on chromVAR scores derived from transcription factor binding (n = 693) and SCP-related chromatin signatures (n = 197). Clones with more than 4 cells were used. Benjamini-Hochberg procedure was used to control false discovery rate. We performed Tukey test to identify clones whose variance of that feature was smaller within the clone than between clones.

### Lineage Barcode Annotation

Only lineage barcodes exceeding a frequency of 0.004 and a clone size of 5 in any *in vitro* sample were included in the analysis. Each lineage barcode was considered to be enriched for an SCP chromatin state or Res 499 enhancer signature if its cells’ z-scored ChromVAR deviation scores were significantly larger than 0 (one-sample t-test). Each lineage barcode was then annotated based on its enriched SCP chromatin states using the following criteria:

1. Lineage barcodes enriched for sensitive SCP signatures and Res 499 deactivated enhancers were annotated as Epi-Sen1/2.
2. Lineage barcodes enriched for resistant SCP signatures and Res 499 activated enhancers were annotated as Epi-Res1 or Epi-Res2.
3. Lineage barcodes enriched for the SCP 11 signature and no other SCP signatures were annotated as Epi-SCP11.
4. Lineage barcodes enriched for both Epi-SCP11 and Epi-Res2 were annotated as hybrid Epi-11/Res2.
5. Lineage barcodes enriched for conflicting SCP signatures were annotated as Plastic.

### STARTRAC pTrans Analysis

STARTRAC^46^ to quantitate developmental relationships between SCP states was run using Watermelon lineage barcodes and SCP chromatin state cell annotations. STARTRAC-pTrans index values were generated for each sample separately to compare how treatment and ICB impacts relationships between states. Bootstrapped 95% confidence intervals were generated by re-sampling the original cells with replacement 1000 times and generating STARTRAC-pTrans index values for each sample. To assess the significance of changes in SCP transitions with JAKi TBK1i treatment, permutation testing was conducted. SCP cell annotations were randomly shuffled, STARTRAC was run on each sample, and the difference between Vehicle and JAKi TBK1i treatment groups was calculated. This process was repeated 1000 times to generate a null distribution. The observed pTrans index difference between Vehicle and JAKi TBK1i treatment groups was compared to the null distribution to generate a P-value.

### Identifying ERE-derived transcripts

We adapted an approach based on *de novo* transcript assembly to identify ERE-derived transcripts^72,73^. We assembled transcripts for each B16 parental, Res 499 parental, and B16 and Res 499 SCP RNA-seq sample separately using StringTie2 (-j 2 -s 5 -f 0.05 -c 2)^74^. We merged all transcripts using TACO version 0.7.3^75^ to generate a unified set of 69,926 assembled transcripts. To narrow down to a set of ERE-derived transcripts, we retained only transcripts that 1. Do not overlap annotated protein-coding exons (GENCODE vM24) and [check that its exons overlapping exons] 2. Are comprised of annotated RepeatMasker repetitive elements in over 20% of exon base pairs. Using this conservative set of criteria, we identified 6,944 high-confidence ERE-derived transcripts. To identify differentially expressed transcripts, we quantified all assembled transcript using featureCounts version 2.0.0^76^ using parameters that distributes multi-mapped reads fractionally across all mapped alignments (-O -M --fraction --primary).

### Identifying ERE Repeat Pairs

We searched for evidence of repeat pairs in ERE-derived transcripts. For each transcript, we identified all overlapping repeat elements (RepeatMasker annotation) and aligned them pairwise in a reverse complement manner using the Smith-Waterman algorithm (Biostrings pairwiseAlignment() function; gap opening and gap extension penalties were 10 and 4, respectively). Pairs that had an alignment score over 25 and were separated by fewer than 6000 base pairs were retained and considered repeat pairs.

